# Agrochemical-responsive gene expression control systems for modulating plant development and metabolism

**DOI:** 10.64898/2026.07.30.741590

**Authors:** Tawni Bull, Lauren Farina, Sam Sutton, Jared Van Blair, Liz Carlsen, Arjun Khakhar

## Abstract

Precise temporal control of gene expression is critical for studying plant biology and engineering complex crop traits. Current systems enable chemically inducible regulation but rely on costly or agriculturally impractical inducers and lack the flexibility needed to regulate combinations of native loci and transgenes. In this work, we elucidate the design rules for control systems, based on Cas9 and Cre recombinase fused to the ecdysone receptor (EcR), which respond to a widely used agrochemical methoxyfenozide (MF). First, we validated the function of both circuits in transient assays and explored how transduction properties can be modulated by engineering nuclear trafficking dynamics. We next characterized both the Cas9-based and recombinase-based systems by using them to regulate fluorescent reporters in transgenic *Arabidopsis thaliana* plants. Here, we demonstrate systemic activation following root application of MF, validating the use of an agriculturally compatible inducer for whole-plant gene regulation. Finally, we validated the utility of the inducible Cas-based SynTF system to regulate multigene pathways and control both metabolic and developmental pathways. Together, these results elucidate design considerations for agrochemical-inducible control systems and demonstrate their utility for temporally regulating plant phenotypes. These synthetic circuits provide a versatile framework for engineering complex traits using an agriculturally compatible inducer.

**Summary:**

- **Research conducted**: Chemically inducible gene expression systems enable temporal regulation of plant traits but are often limited by costly, agriculturally impractical inducers and inflexible architectures. We developed methoxyfenozide (MF)-responsive Cas9-based synthetic transcription factor (SynTF) and Cre recombinase control systems to enable programmable regulation of endogenous genes and transgenes.
- **Methods:** Cas9- and Cre-based circuits fused to the ecdysone receptor (EcR) were engineered and characterized in transient *Nicotiana benthamiana* assays and stable *Arabidopsis thaliana* transgenic lines. We evaluated the effects of engineering nuclear trafficking dynamics and tested inducible regulation of reporter genes, metabolic pathways, and plant development.
- **Key results:** Both control systems enabled MF-dependent reporter gene activation in transient assays and in stable lines via systemic whole-plant induction following root application of MF. Engineering nuclear trafficking improved system performance by modulating leakiness and dynamic range. The Cas9-based control system enabled regulation of multigene metabolic and developmental pathways, demonstrating programmable temporal control of complex traits.
- **Main conclusion:** These results establish design principles for agrochemical-inducible genetic control systems and demonstrate that agriculturally compatible inducers can provide flexible, temporal regulation of plant metabolism and development, creating a versatile platform for engineering complex plant traits.

## Introduction

Control systems that modulate gene expression in response to chemical inducers are useful for fundamental plant biology and crop trait engineering (1–7). By enabling temporal regulation and dose-dependent tuning of gene expression, these systems provide a valuable approach for engineering traits associated with trade-offs, such as organ size or pathogen resistance (8–11). For agricultural deployment, inducers must be inexpensive, field stable, and non-toxic. Agrochemicals, including commonly used insecticides, meet these criteria and are formulated for efficient uptake and systemic transport through vascular tissues (12, 13). Therefore, agrochemical-inducible control systems could enable gene regulation throughout the plant, including internal tissues that are challenging to reach with topical application. Ideal systems should also exhibit tight off states, high induction ratios, compact genetic architectures, and the flexibility to regulate both native genes and transgenes (14).

Prior studies have identified signaling domains that sense commonly used agrochemicals to trigger DNA binding, protein-protein interactions, or nuclear localization in plants (1, 15). However, these systems often require coupling with secondary control systems that are more flexibly programable, such as Cas9-based synthetic transcription factors (SynTFs) (1), to regulate genes not driven by their target synthetic promoters. This significantly increases the size of genetic cargos and can introduce other suboptimalities, such as delayed responses to inducers, leaky off states, and a lack of dose dependency (16–18). Thus, strategies are needed to integrate agrochemical-sensing domains with flexibly programable regulators to generate compact control systems.

Among existing inducible systems in plants, the dexamethasone-inducible SynTF has been widely used for basic science due to its tight off state, high fold change, and dose dependent activation (2, 6, 19). It consists of a nuclear receptor fused to DNA-binding and transcriptional activation domains, enabling ligand-gated nuclear localization and subsequent activation of a synthetic promoter. Although widely used in research, dexamethasone-inducible systems are not suited for agricultural deployment because dexamethasone is expensive, toxic in high doses (20), and not approved for agricultural use. In addition, it has been shown to cause growth defects in transgenic plants (21).

To overcome this challenge, prior studies developed chemically inducible plant controls systems based on the non-steroidal ecdysone agonist methoxyfenozide (MF), the active ingredient in the commercially available insecticide, Intrepid2F (4, 5, 22). These systems fuse the ligand binding domain (LBD) of the spruce budworm (*Choristoneura fumiferana*) ecdysone receptor to a DNA binding domain and a transcriptional activation domain (4). Upon ligand binding, the fusion protein accumulates in the nucleus, binds *cis*-regulatory elements of a synthetic promoter, and activates gene expression. Initial studies demonstrated robust luciferase induction in transgenic *Arabidopsis* and tobacco with minimal background expression (4). Subsequent mutagenesis of the ecdysone LBD further improved MF sensitivity, enabling stronger induction at lower ligand concentrations (23).

Taken together, these studies establish ecdysone-based systems as effective control systems for inducible transgene expression in plants. However, whether the nuclear trafficking dynamics of such control systems can be engineered to improve transduction properties remains unexplored. Furthermore, existing ecdysone-based control systems regulate transgenes through synthetic promoters and have not been integrated with programmable genetic control systems, such as Cas9-based SynTFs and site-specific recombinases. Integrating the ecdysone LBD with these tools would generate compact control systems for inducible regulation of endogenous genes and genetic state transitions, thereby enabling sophisticated control of crop traits.

In this work we elucidate design considerations for integrating the ecdysone LBD signaling domain with Cas9- and Cre recombinase-based genetic control systems to enable programmable, MF-inducible modulation of gene expression. We first characterize system performance in transient *Nicotiana benthamiana* agroinfiltration assays and investigate whether ligand-gated nuclear trafficking can enhance their signaling properties. We then evaluate system function in stable *Arabidopsis thaliana* transgenic lines using a fluorescent reporter. Building on this, we demonstrate inducible transcriptional modulation of an endogenous metabolic pathway to enhance luminescence output from a multigene synthetic metabolic pathway. Finally, we explore our system’s utility for inducible control of plant development while mitigating pleiotropic effects. Together, these studies establish a framework for programmable and chemical control of gene regulation and highlight its potential for future crop trait engineering.

## Materials and Methods

### Plasmid construction

All plasmids constructed in this work were built using Golden Gate Assembly and Modular cloning (24–26) and are listed with links to their annotated plasmid maps in **Supplemental Table S1**. DNA parts were amplified from previously reported plasmids (24, 27–31) or synthesized using Twist Bioscience. Guide RNA (gRNA) target sites were designed using CHOPCHOP v3 (32) and the target sequences are listed in **Supplemental Table S2.** More detailed methods on plasmid construction are in **Supplemental Information**.

### Transient assays

Cultures of *Agrobacterium tumefaciens* strain GV3101 carrying the SynTF and/or reporter plasmids were prepared and infiltrated using a previously published protocol (33) with few modifications. One day after infiltration, plants were watered with 500 mL of 2.2 mM methoxyfenozide as Intrepid 2F (MF) or tap water. Two days post induction (DPI), leaves were samples and fluorescence was quantified as described below. More detailed methods on transient assays are in **Supplemental Information**.

### Stable line generation

All stable *Arabidopsis thaliana* transgenic lines in this work were generated using floral dip (34). For the Cas9-EcR control system, the Cas9-EcR T-DNA plasmids were first transformed into *A. thaliana* followed by transformation with the ratiometric fluorescent reporter with and without the gRNAs targeting mScarlet for activation, *DELLAs* for activation, or the FBP with and without gRNAs targeting the phenylpropanoid pathway. T1 plants were used for further experiments. Cre recombinase T-DNA plasmids with and without the reporter were transformed into *A. thaliana* and T1 plants were used for further experiments. More detailed methods for the generation of transgenics are available in **Supplemental Information**.

### Methoxyfenozide induction of stable transgenic lines

Seeds of EcR-Cre, EcR-Cre HIV1 REV, and *DELLA* activation independent transgenic lines were sterilized and plated as described previously (28). Ten-day old seedlings were transferred to 1/2x MS agar plates supplemented with 80nM or DMSO as a negative control. Fluorescence and root length measurements were quantified at multiple timepoints as described in **Supplemental Information**.

Seeds for all the *A. thaliana* lines containing Cas9-EcR were sterilized and plated as described previously (28). Cas9-EcR, Cre recombinase, and *DELLA* activation lines were germinated on 1/2x MS agar plates supplemented with appropriate antibiotics. For the demethylation experiment, plates also contained 50uM 5-Azacitidine (AzaC). For all experiments, 14-day old seedlings were transplanted into soil. After two weeks, a subset of plants was top watered twice per week with either 31 mL of 2.2 mM MF, 0.5 mM MF, or tap water. After 14 days of induction (4 total applications), fluorescence, luminescence, or shoot phenotypes were quantified. More detailed methods are available in **Supplemental Information**.

### Quantification of reporter fluorescence and luminescence output

For all transient and soil assays, reporter fluorescence was quantified using a Tecan Spark plate reader. Leaf discs were harvested from infiltration sites of *N. benthamiana* transient assays or rosette leaves from *A. thaliana* stable lines and Venus and mScarlet measurements were performed. For methoxyfenozide plate assays, mScarlet fluorescence was quantified using an Analytik Jena UVP ChemStudio PLUS Imaging System. Image acquisition settings were held constant across all samples. Fluorescence intensity was quantified in Fiji (ImageJ2). More detailed information on fluorescence quantification for each experiment is available in **Supplemental Information**.

Reporter luminescence was quantified using a AnalytikJena UVP ChemStudio PLUS Imaging System and VisionWorks imaging software. Image acquisition settings were held constant across all samples and resulting images were analyzed using the VisionWorks Area Density tool. More detailed information on luminescence quantification is available in **Supplemental Information**.

### *Arabidopsis* shoot and germination phenotyping

Phenotyping of *A. thaliana* shoots was performed at 14 DPI. Individual plants were photographed using a Canon EOS Rebel T6 camera and rosette area estimates were measured in Fiji by tracing the outer perimeter of the rosette. For leaf areas, plants were dissected by leaf, scanned, and measured in Fiji.

For evaluating germination efficiency of the *DELLA* activation lines, independent lines of the Cas9-EcR *DELLA* activation and constitutive *DELLA* activation lines were sown on 1/2x MS agar plates as described previously (28). The total number of germinated seeds was collected at multiple timepoints, and the germination percentage of each genotype was calculated. More detailed information on phenotyping is available in **Supplemental Information**.

### Gene expression analysis

The relative expression of target genes was measured using a modified version of a previously reported protocol (28, 35). Concentrations of all genes were quantified using RT-qPCR performed using Luna® Universal qPCR Master Mix on a BioRad CFX Opus qPCR machine. Cycle threshold values for all genes were normalized to the housekeeping gene, *PP2A*. Gene specific, PCR, and RT-qPCR primers are listed in **Supplemental Table S4**. More details on gene expression analysis methods are available in **Supplemental Information**.

### Data analysis

All data analysis was performed in Python using Jupyter Lab (version 3.4.4). More information on data analysis is available in **Supplemental Information**. All the raw data and code used to analyze the data is available on the following github repository: https://github.com/arjunkhakhar/260729_Methoxyfenozide_Control_Systems.

## Results

### Agrochemical-responsive control systems enable inducible activation of reporters in transient expression assays

Previous work demonstrated the programmability of Cas9-based SynTFs for regulating gene expression in plants (28, 29). These SynTFs contain constitutively expressed, nuclear-localized Cas9, an RNA-binding protein (RBP) fused to the transcriptional trans-activator DREB2A (MCP-DREB2A), and an RBP fused to a truncated transcriptional repressor, TOPLESS (PCP-TPLN300) (28). A guide RNA (gRNA) truncated to 14 base pairs and modified with a 3’ MS2 or PP7 RBP-recruitment motif (a gRNA scaffold) enables Cas9 binding without DNA cleavage, allowing SynTF assembly at the target promoter (36). To enable ligand-inducible regulation, we replaced the C-terminal nuclear localization sequence on Cas9 with a previously validated mutated ecdysone receptor (EcR) that is highly sensitive to MF formulated as Intrepid2F (37). Upon Intrepid2F treatment, Cas9-EcR is expected to translocate to the nucleus, enabling SynTF assembly and target gene regulation (**Fig. 1A**).

**Figure 1.**
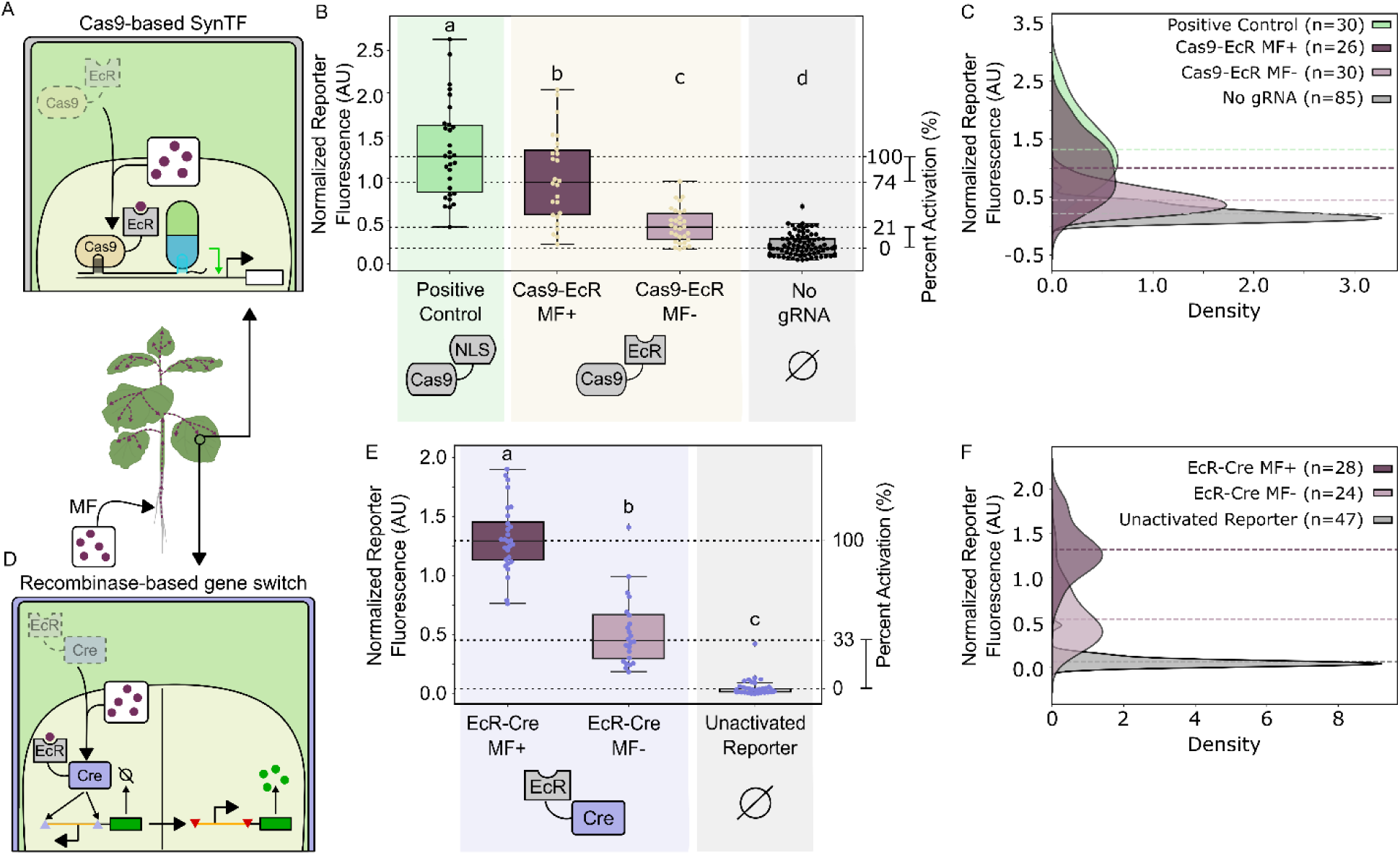
Transient validation of the CasG-and Cre-recombinase agrochemical inducible control systems. A) Schemic depicting the function of the EcR-Cas9-based control system tested in transient agroinfiltration assays in *N. benthamiana*. B) Boxplots representing the normalized reporter fluorescence from transient assays of plants co-infiltrated with the Cas9-based SynTF and treated with (dark magenta, MF +, n=9 leaves across 4 independent plants) and without (light magenta, MF -, n=10 leaves across 4 independent plants) MF. The Cas9-EcR system is compared to an Cas9-NLS as a positive control (green, n=10 leaves across 4 independent plants) and the Cas9-EcR control system without gRNAs (gray, n=19 leaves across 8 independent plants) as a negative control. C) Kernal density estimation (KDE) plots representing the distributions of the normalized reporter fluorescence for the Cas9-based control system. The horizontal dashed lines represent the mean normalized reporter fluorescence for each system. The total number of fluorescence measurements (leaf discs) is stated in the figure legend. D) Schematic depicting the function of the Cre recombinase-based inducible control system tested in transient agroinfiltration assays in *N. benthamiana*. E) Boxplots representing the normalized reporter fluorescence from transient assays of plants infiltrated with the EcR-Cre control system and treated with (dark magenta, MF +, n=10 leaves across 3 independent plants) and without (light magenta, MF -, n=11 leaves across 3 independent plants) MF compared to an unactivated reporter (gray, n=20 leaves across 6 independent plants). F) KDE plots representing the distributions of normalized reporter fluorescence of the Cre-based control system. The horizontal dashed lines represent the mean normalized reporter fluorescence for each system. The total number of fluorescence measurements (leaf discs) is stated in the figure legend. Across all plots, every data point represents a normalized fluorescence measurement from a single leaf disc. Different letters represent statistically significant differences (One-way ANOVA followed by Tukey HSD test, *p* < 0.05).

To test the functionality of the system, we performed transient agroinfiltration assays in *Nicotiana benthamiana* using a ratiometric reporter (**Figure S1**). The SynTF was targeted to a synthetic promoter consisting of four tandem repeats of the Lac operon followed by the minimal CaMV35S promoter (4xLacO pMin35s), which drove expression of the reporter gene, mScarlet. A constitutively expressed Venus reporter was included for normalization of differences in T-DNA delivery (**Figure S1A**). An *A. tumefaciens* strains harboring the SynTF T-DNA plasmid the reporter T-DNA plasmid encoding expression cassettes for both fluorescent proteins and a gRNA scaffold targeted to the LacO elements were co-infiltrated into *N. benthamiana* (**Figure S1A**). As a negative control, the SynTF strain was co-infiltrated with a reporter plasmid lacking the gRNA scaffold. As a positive control, the SynTF strain contained a constitutive Cas9-NLS, rather than Cas9-EcR. One day after infiltration, plants were water with or without MF (**Figure S1B, S1C**). Two days post induction (DPI), changes in reporter fluorescence were quantified. Plants co-infiltrated with gRNAs and treated with MF resulted in a 2.23-fold increase in mean normalized fluorescence relative to the untreated controls (*p* = 1.41 x 10^-8^, **Figure1B**), demonstrating successful ligand-dependent activation. However, we observed only 74% of the normalized fluorescence in the treated Cas9-EcR plants (*p* = 0.0024) when compared to the Cas9-NLS positive control (**Figure 1B**, **1C**), indicating incomplete activation of the system. Furthermore, untreated Cas9-EcR samples displayed significantly higher mean normalized fluorescence than the no gRNA negative controls (2.07-fold increase, *p* = 0.0057, **Figure 1B**, **1C**), revealing a significant leak of the Cas9-EcR system, even in the absence of the inducer. Consistent with these findings, the normalized fluorescence distributions of the treated and untreated samples showed considerable overlap (0.407 Gaussian KDE overlap coefficient, **Figure 1C**). Together, these results demonstrate that integrating EcR with Cas9 enables programmable agrochemical inducible gene activation. However, both incomplete activation in the “on” state and leaky activation in the “off” necessitate improvement of the transduction properties of the system.

While the Cas9-EcR inducible SynTF control system enables programable and reversible induction of expression, the reliance on continuous ligand application limits its utility in certain contexts. In contrast, irreversible activation of transgene expression can be advantageous in contexts where a stable or self-maintaining expression state is desired. To address this limitation, we integrated EcR with a Cre recombinase-based control system (38–41) that irreversibly flips a promoter from the antisense direction to the sense direction via site specific recombination (**Figure 1D**). Here, we fused EcR to the N-terminus of Cre (EcR-Cre) and built a ratiometric reporter T-DNA plasmid. The regulated reporter consisted of the *A. thaliana* UBIQUITIN10 promoter in the antisense orientation and flanked by Lox sites, which are targeted by the recombinase and are upstream of the mScarlet reporter. The ratiometric reporter also included the same constitutively expressed Venus reporter for normalization as described above. In the basal (“off”) state, the promoter driving expression of mScarlet is in the antisense orientation preventing transcription. Following MF application, ligand-dependent accumulation of the EcR–Cre fusion protein in the nucleus facilitates recombination at the flanking Lox sites. This recombination event inverts the promoter into the sense orientation which breaks the Lox sites, leading to irreversible activation of transgene expression (**Figure 1D**).

We tested the inducibility of the EcR-Cre control system using agroinfiltration assays in *N. benthamiana* as described previously. Two days after application of MF, we observed a significant increase in mean normalized fluorescence of treated samples compared to the untreated controls (1.91-fold increase, *p* = 7.32 x 10^-10^), **Figure 1E**). Further experiments exhibited EcR-Cre activation levels similar to a nuclear localized Cre, demonstrating complete activation of the system (**Figure S2**). However, similar to the Cas9-EcR control system, we also observed leaky activation of the EcR-Cre control system in the absence of the inducer. This was evident from the higher mean normalized fluorescence of the untreated samples relative to the negative control treatment (*p* = 6.63 x 10^-6^, **Figure 1E**). The magnitude of leak observed when comparing the untreated controls to the treated samples with the EcR-Cre system was larger (52% activation in the ‘off’ state, **Figure 1E**) than the leak observed with the Cas9-EcR system (45% activation in the ‘off’ state, **Figure1B**). This is likely the result of irreversible activation enabling leak to accumulate over time. Although the magnitude of leak was higher in the EcR-Cre system, we also observed less overlap between the treated and untreated samples (0.225 Gaussian KDE overlap coefficient, **Figure 1F**). This is consistent with the expected behavior of a binary genetic switch, in which activation results in discrete on and off states rather than a graded response.

### Engineering nuclear trafficking of EcR-based control systems reduces leak

To reduce leak, we fused a nuclear export sequence (NES) to the C-termini of Cas9-EcR (Cas9-EcR-NES) and EcR-Cre (EcR-Cre-NES). We hypothesized this would promote cytoplasmic accumulation in the absence of the inducer, thereby reducing leak in the off state (**Figure 2A**, **2C**). We tested two previously described NES variants: the 23 amino acid motif from the squash leaf curl virus movement protein, BR1 (BR1 residues 177-199) and the 9 amino acid motif from the human immunodeficiency virus type 1 rev protein (HIV-1 REV residues 75-83) (42).

**Figure 2.**
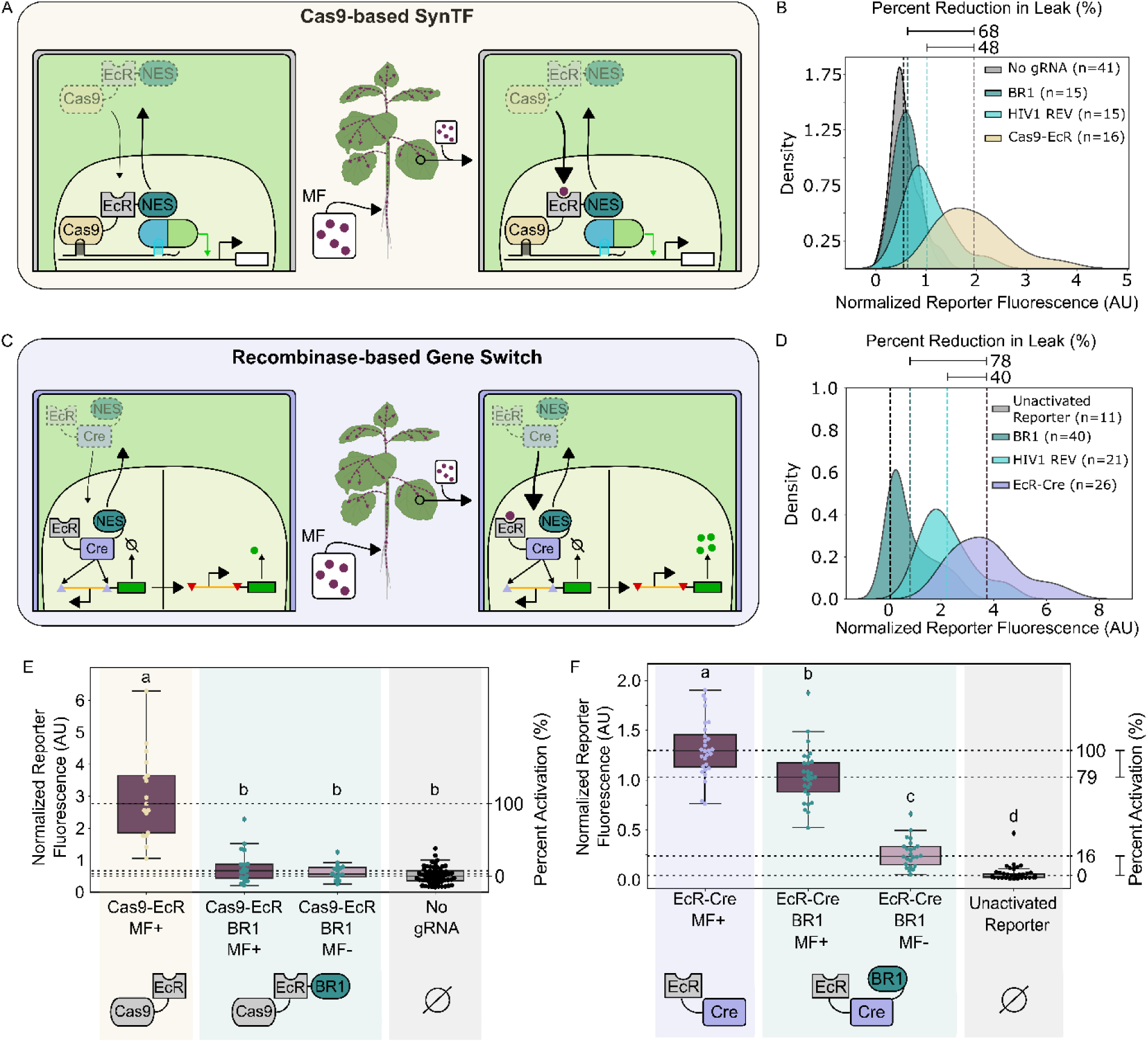
Engineering nuclear trafficking dynamics to improve the transduction of the CasG-and Cre recombinase-based agrochemical inducible control systems. A) Schematic depicting the incorporation of an NES to the Cas9-based SynTF inducible control system tested in transient agroinfiltration assays in *N. benthamiana*. B) Kernal density estimation (KDE) plots representing the distributions of the normalized reporter fluorescence for the Cas9-EcR (tan), Cas9-EcR-HIV1 REV NES (light teal), and Cas9-EcR-BR1 NES (dark teal) in the off state. The vertical dashed lines represent the mean normalized reporter fluorescence for each version of the system. The total number of fluorescence measurements (leaf discs) is stated in the figure legend. C) Schematic depicting the incorporation of an NES into the recombinase-based inducible control system tested in transient agroinfiltration assays in *N. benthamiana*. D) KDE plots representing the distributions of the normalized reporter fluorescence for the EcR-Cre (purple), EcR-Cre-HIV1 REV NES (light teal), and EcR-Cre-BR1 NES (dark teal) in the off state. The vertical dashed lines represent the mean normalized reporter fluorescence for each version of the system. The total number of fluorescence measurements (leaf discs) is stated in the figure legend. E) Boxplots representing the normalized reporter fluorescence of plants co-infiltrated with the Cas9-EcR control systems and treated with (dark magenta, MF +, n=6 leaves across 3 independent plants) or without (light magenta, MF -, n=6 leaves across 3 independent plants) MF. F) Boxplots representing the normalized reporter fluorescence of plants infiltrated with the EcR-Cre control systems and treated with (dark magenta, MF +, n=10 leaves across 3 independent plants) or without (light magenta, MF -, n=10 leaves across 3 independent plants) MF. Across all plots, every data point represents a normalized fluorescence measurement from a single leaf disc. Different letters represent statistically significant differences (One-way ANOVA followed by Tukey HSD test, *p* < 0.05).

We first tested the impact of each NES on the inducible Cas9-based SynTF. We repeated the transient agroinfiltration assays in *N. benthamiana* as described above, wherein Cas9-EcR-NES strains were co-infiltrated with the same ratiometric reporter used previously. When either BR1 (Cas9-EcR-BR1) or HIV1 REV (Cas9-EcR-HIV1 REV) NES was fused to the Cas9-EcR control system, we observed a significant reduction in basal leak relative to the unmodified Cas9-EcR, with BR1 and HIV-1 REV decreasing leak by 68% (*p* = 1.48 x 10^-11^) and 48% (*p* = 9.58 x 10^-7^), respectively (**Figure 2B**). In both cases, the mean normalized fluorescence was not significantly different from the negative controls (**Figure 2E, S3A**), indicating effective suppression of off-state activity. However, both NES fusions also abolished inducibility as MF treatment failed to produce a significant increase in reporter induction relative to uninduced controls (**Figure 2E, S3A**). This resulted in a lower fold change in induction for each of the NES fusions when compared to Cas9-EcR (**Figure S3B**). These results suggest that incorporation of an NES leads to excessive cytoplasmic retention of the SynTF, preventing sufficient nuclear accumulation for efficient regulation upon ligand activation.

We next applied the same nuclear export strategy to the recombinase-based system by fusing either the BR1 (EcR-Cre-BR1) or HIV1 REV (EcR-Cre-HIV1 REV) NES to the C-terminus of EcR-Cre and repeated the transient agroinfiltration assay as described above. Similar to Cas9-based system, NES incorporation improved the performance of the recombinase-based control system by significantly reducing the leak compared to the unmodified EcR-Cre by 78% (*p* = 1.24 x 10^-12^) and 40% (*p* = 3.71 x 10^-12^) for BR1 and HIV REV1, respectively (**Figure 2D**). In contrast to the Cas9-based system, induction of reporter fluorescence for each NES variant was maintained with activation levels at 79% and 83% of the unmodified EcR-Cre control system for BR1 and HIV1 REV, respectively (**Figure 2F, S4A**). As a result, the dynamic range of the system significantly increased for both NES variants relative to the unmodified control system, where BR1 led to a larger improvement in dynamic range (1.59-fold increase, *p* = 9.02 x 10^-8^) than HIV1 REV (1.46-fold increase, *p* = 0.008; **Figure S4B, S4C**). This is consistent with our results characterizing the Cas9-EcR systems fused to these NES sequences, wherein HIV1 REV confers weaker nuclear export than BR1.

### Agrochemical-responsive recombinases enable induction of reporter expression in transgenic plants

Thus far, transient assays have demonstrated inducible reporter gene expression, however, these are limited by localized expression in leaf tissue and do not establish whole-plant performance. We therefore generated *Arabidopsis thaliana* transgenic lines expressing the recombinase-based (EcR-Cre, EcR-Cre-HIV1 REV, and EcR-Cre-BR1) systems together with the ratiometric reporter described above, as they exhibited the highest dynamic range in transient assays.

We screened independent *A. thaliana* lines for MF responsiveness. EcR-Cre and EcR-Cre-HIV1 REV seedlings were grown on agar plates for 10 days, then transferred to plates supplemented with 80 nM pure MF or control plates. Whole-plate fluorescence images were acquired over three days, and mScarlet fluorescence was quantified for each seedling (**Figure 3A**, **3B, S5**). To account for basal leak, fluorescence at 24 and 72 hours was corrected by subtracting each seedling’s pre-induction (0 hour) value.

**Figure 3.**
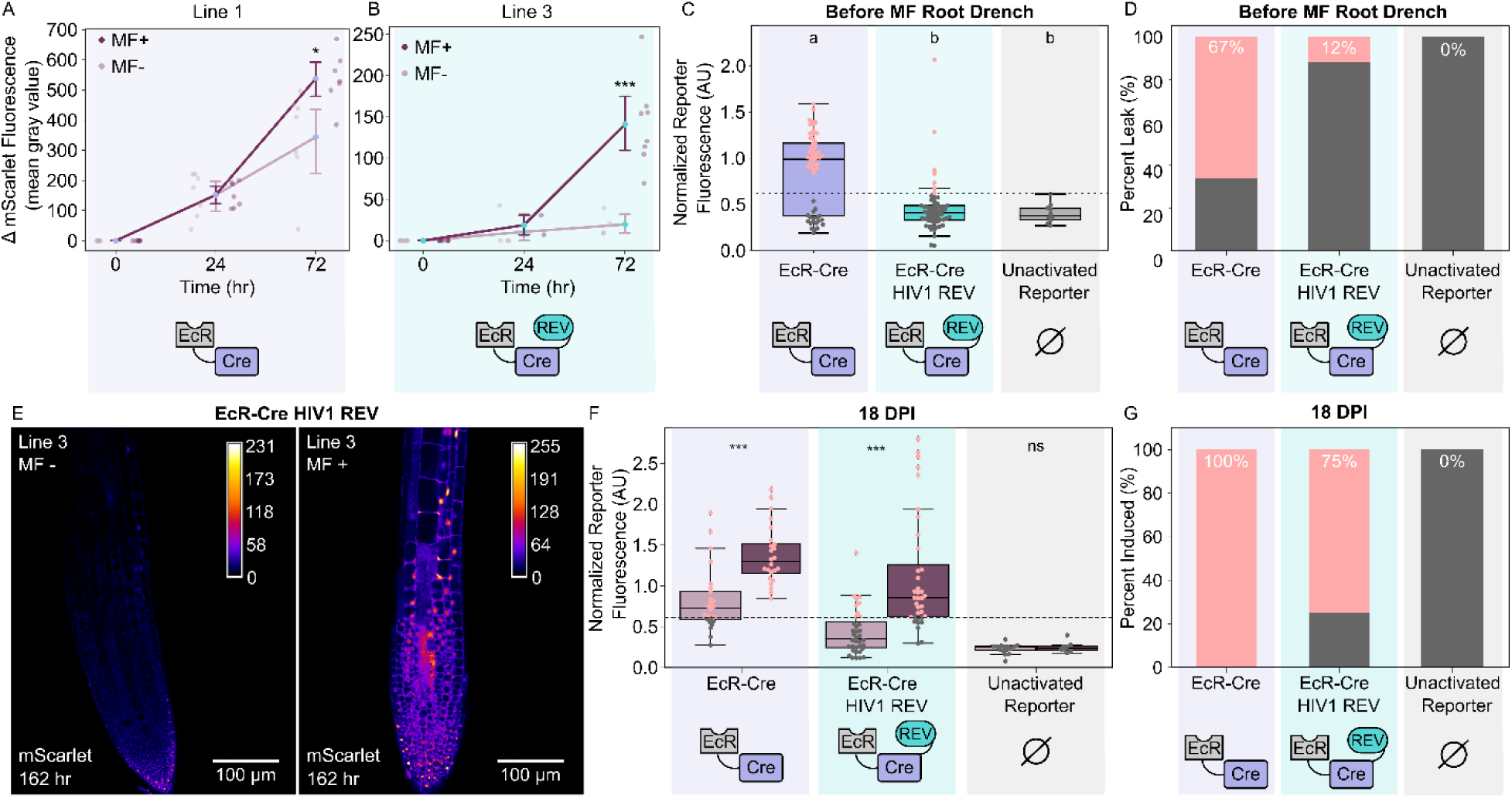
Characterization of the Cre-recombinase control system in stable transgenic lines. A, B) Line plots representing the mScarlet fluorescence over 72 hours for an EcR-Cre (A) or EcR-Cre HIV1 REV (B) *A. thaliana* independent transgenic line treated with (dark magenta, MF +) or without (light magenta, MF -) MF. Each data point represents the fluorescent measurement from an individual plant. C) Boxplot representing the normalized reporter fluorescence of leaf samples collected from EcR-Cre (n=8 plants each from 2 independent lines), EcR-Cre HIV1 REV1 (n=8 plants each from 3 independent lines), and an unactivated reporter population (n=10 plants from 1 independent line) before treatment with MF. The horizontal dashed line represents the leak threshold. Pink colored dots represent samples that exceeded the leak threshold for each genotype. D) Stacked barplot depicting the percentage of leaf discs displaying leak for EcR-Cre (n=57 measurements across 2 independent lines), EcR-Cre-HIV1 REV (n=60 measurements across 3 independent lines), and an unactivated reporter (n=10 measurements from 1 independent line). E) Representative confocal microscopy images of the root tip from an EcR-Cre HIV1 REV plant (Line 3) treated with (right) or without (left) MF. The heat map represents the intensity of mScarlet fluorescence. F) Boxplots representing the normalized reporter fluorescence of independent lines of EcR-Cre (n=4 plants across 2 independent lines for both MF+ and MF-treatments), EcR-Cre HIV1 REV (n=4 plants across 3 independent lines for both MF+ and MF-treatments), and an unactivated reporter population (n=3 or 5 plants from 1 independent line for MF+ and MF-treatments, respectively) treated with (MF+, dark magenta) or without (MF-, light magenta) MF. G) Stacked barplots depicting the percentage of leaf discs displaying an induced phenotype after treatment with MF for EcR-Cre (n=24 measurements across 2 independent lines), EcR-Cre-HIV1 REV (36 measurements across 3 independent lines), and an unactivated reporter population (8 measurements from 1 independent line). Across all plots, every data point represents a normalized fluorescence measurement from a single leaf disc. Different letters represent statistically significant differences (One-way ANOVA followed by Tukey HSD test, *p* < 0.05). Asterisks represent results from a Welch’s two sample *t*-test (*p* < 0.05), * corresponds to *p* < 0.05, ** corresponds to *p* < 0.005, and *** corresponds to *p* < 0.0005.

After 72 hours, MF-treated seedlings exhibited significantly greater mScarlet fluorescence than untreated controls in both systems (**Figure 3A**, **3B**). The best performing EcR-Cre line (Line 1) showed a 1.57-fold increase in fluorescence relative to the untreated controls (*p* = 0.018, **Figure 3A**). Line 2 also showed a 1.28-fold increase, although this difference was not significant (**Figure S5A**). In the EcR-Cre-HIV1 REV system, Line 2 exhibited a 2.09-fold increase (*p* = 0.025, **Figure S5B**), while Line 3 showed the strongest response, with a 7.19-fold increase (*p* = 0.0002, **Figure 3B**, **3E, S6**). Both recombinase systems therefore remained functional in stable lines, with the EcR-Cre-HIV1 REV system showing greater dynamic range than EcR-Cre, consistent with transient assays (**Figure S4B**, **S4C**).

We next assessed basal leakiness of the recombinase-based control system in soil-grown plants by screening leaves for reporter activation before MF treatment. The EcR-Cre system exhibited a bimodal distribution of normalized reporter fluorescence, with 67% of sampled leaves exceeding the negative control threshold, defined as the highest normalized fluorescent value observed in the unactivated reporter (**Figure 3C**, **3D**). In contrast, transgenic lines harboring the EcR-Cre-HIV1 REV control system exhibited lower background activity, with only 12% of sampled leaves displaying leak above the negative control threshold (**Figure 3C**, **3D**). These findings agree with transient assays, in which incorporation of an NES resulted in a reduction of leak in the system.

To test the systemic response of our control system, half of the soil-grown plants were treated with an MF root drench (Intrepid2F) and the remaining plants with water. Fluorescence measurements were then collected from leaves of both treatments. At 18 DPI, both the EcR-Cre and the EcR-Cre-HIV1 REV systems exhibited significantly increased mean normalized fluorescence (mScarlet/Venus) compared to their respective uninduced controls (**Figure 3F, S5C**). Consistent with the transient assays, the EcR-Cre-HIV1 REV system exhibited lower basal leakiness in the uninduced state while retaining its inducibility, with 75% of MF-treated leaves exceeding the negative control threshold defined previously (**Figure 3G**). Although this proportion was lower than that of the unmodified EcR-Cre system, in which 100% of MF-treated leaves exceeded the negative control threshold, the EcR-Cre-HIV1 REV control system exhibited a significantly greater fold induction (2.93-fold increase) owing to its reduced background leakiness and improved dynamic range **(Figure 3F**, **3G, S5C**). Like transient assays and consistent with previous studies, these results demonstrate that MF applied via a root drench can induce recombinase-mediated gene activation across the plant body.

Interestingly, the EcR-Cre-BR1 control system consistently produced the weakest induction among the three versions tested (**Figure S7**). This finding differs from the observations in transient assays, in which the BR1 NES resulted in the largest dynamic range (**Figure S4B**). This discrepancy may reflect differences in NES activity across *N. benthamiana* and *A. thaliana*.

### Silencing can inhibit the function of recombinase-based control systems

Despite successful induction during initial experiments, repeated plate-and soil-based induction assays using the same Cre recombinase-based stable lines failed to produce significant increases in normalized reporter fluorescence. Previous studies have reported that loxP-containing expression cassettes can become hypermethylated in plants, resulting in epigenetic silencing of transgenes (43). We, therefore, hypothesized that methylation-mediated silencing of the reporter cassette reduced inducibility over time. To test this, we collected genomic DNA from bulked samples of representative EcR-Cre (Line 1) and EcR-Cre-HIV1 REV (Line 3) lines treated with MF and used PCR to screen for the presence of the promoter in the sense orientation. In both control systems, we observed the presence of the recombined promoter while simultaneously observing no significant increase in mScarlet fluorescence after 72 hours (**Figure S8**). This data provides evidence that recombination is occurring even with the lack of fluorescence signal suggesting that silencing of gene expression may be a contributing factor to the loss of signal overtime.

To further test if methylation-mediated silencing contributed to the loss of signal, plants of the same EcR-Cre (Line 1) and EcR-Cre-HIV1 REV (Line 3) lines were treated with the DNA demethylating agent, 5-azacytidine (azaC) (43). We hypothesized that if methylation was responsible for the loss of inducibility, azaC should restore reporter activation following MF application. Plants were germinated on agar plates supplemented with azaC, established in soil, induced with an MF root drench, and screened for reporter fluorescence. Consistent with our hypothesis, azaC treatment partially restored inducible reporter expression in both versions of the control system (**Figure SG**). At 14 DPI, treated plants of both EcR-Cre and EcR-Cre-HIV1 REV lines showed significantly increased mean normalized reporter fluorescence relative to their corresponding untreated controls (**Figure SG**). Although the magnitude of induction was lower than that observed during the initial screen, both EcR-Cre (1.61-fold increase, *p* = 0.02) and EcR-Cre-HIV1 REV (1.20-fold increase, *p* = 0.001) exhibited significant recovery of inducibility (**Figure SG**). These findings provide evidence that DNA methylation contributes to silencing of the loxP-containing reporter cassette and limits long-term performance of control systems that rely on this recombinase.

### Agrochemical-inducible CasG-based SynTFs enable temporal and dose-dependent control of expression

Due to the transgene silencing limitations of the recombinase-based system and the need for programmable regulation in some contexts, we further characterized the Cas9-EcR system in whole plants. When engineering aspects of development or metabolism that are controlled by multiple genes and associated with trade-offs, the capacity to control the timing or strength of expression of multiple genes is valuable (8, 14, 44). In this context, inducible control systems that enable reversible and programmable regulation may help mitigate these trade-offs. Our Cas9-EcR control system provides an ideal framework for dynamic and multiplexed regulation of complex pathways.

We evaluated the performance of our Cas9-EcR system in stable lines by building *A. thaliana* plants that co-express the Cas9-EcR SynTF with the ratiometric reporter described above (**Figure 4A**). As a control, we generated additional transgenic lines that harbored the SynTF and the reporter but lacked the gRNAs (no gRNA control). An initial screen for inducible activation in whole plants identified three independent lines that exhibited significant reporter activation under treated conditions when compared to their respective untreated controls. To quantify this response, we calculated the fold change in mean normalized reporter fluorescence (induced sample/mean of the uninduced samples) for each independent line as well as the no gRNA control population. At 14 DPI, all three lines resulted in a significant increase in reporter fluorescence when compared to the control lines (**Figure 4B**).

**Figure 4.**
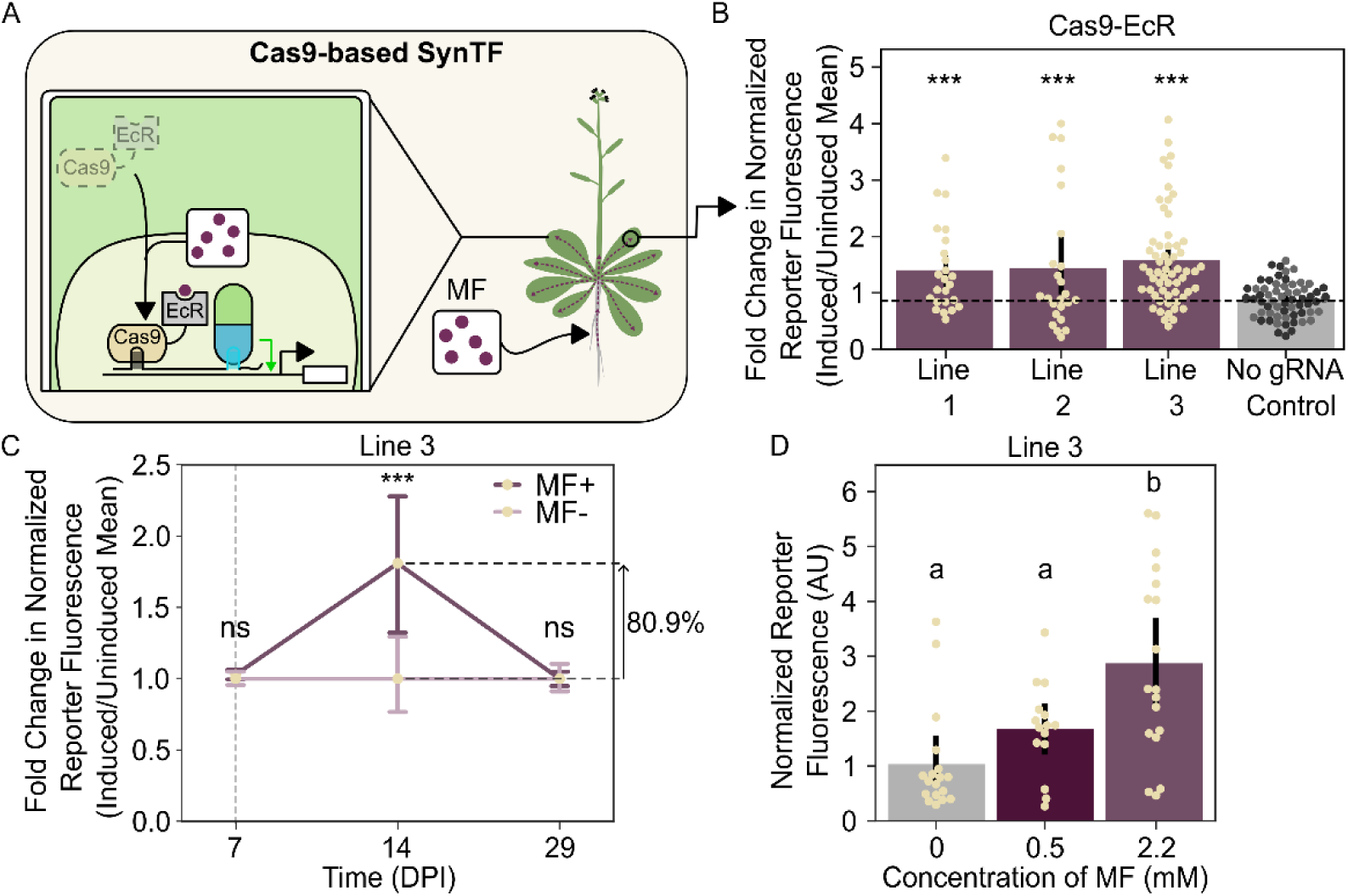
Characterization of the CasG-based control system in stable transgenic lines. A) Schematic depicting the Cas9-based SynTF inducible control system. B) Barplot representing the fold change in normalized reporter fluorescence (induced/ mean of uninduced samples) of three independent lines (n=8 plants per independent line per treatment) compared to a population of no gRNA control lines (8 plants across 2 independent lines per treatment). Every data point represents a normalized fluorescence measurement from a single leaf disc. C) Line plot representing the fold change in normalized reporter fluorescence (induced/mean of uninduced samples) of leaf discs over time from a population of plants (Line 3, n=8 plants per treatment) treated with (dark magenta, MF +) and without (light magenta, MF -) MF. The gray dashed line represents when MF treatment stopped. D) Barplots depicting the normalized reporter fluorescence of leaf discs collected from plants (Line 3) treated with 0 (n=15 leaves across 5 plants), 0.5 (n=12 leaves across 4 plants), or 2.2 mM (n=15 leaves across 5 plants) MF at 14 DPI. Every data point represents a normalized fluorescence measurement from a single leaf disc. Asterisks represent results from a Welch’s two sample *t*-test (*p* < 0.05), * corresponds to *p* < 0.05, ** corresponds to *p* < 0.005, and *** corresponds to *p* < 0.0005. Different letters represent statistically significant differences (One-way ANOVA followed by Tukey HSD test, *p* < 0.05).

To further characterize the induction dynamics in stable plants, we next focused on Line 3, which displayed the strongest response to MF application. Soil grown plants were treated with and without MF and reporter expression was quantified at multiple timepoints.

At 7 DPI, Line 3 showed a moderate increase (2.8%) in mean normalized reporter signal under induced conditions, which was not statistically significant when compared to uninduced plants (**Figure 4C**). By 14 DPI, signal increased substantially, displaying a 80.9% increase in mean normalized reporter fluorescence when compared to uninduced plants (*p* = 6.69 x 10^-3^, **Figure 4C**). At 29 DPI, 22 days after the final root drench, reporter fluorescence had returned to levels comparable to those observed in the no gRNA population (*p* = 0.981, **Figure 4C**), demonstrating reversibility of the induced response.

Interestingly, we also observed that baseline normalized signal in the no gRNA control plants decreased significantly following MF treatment (11.9% decrease, *p* = 0.024, **Figure S10A**), suggesting the agrochemical reduces reporter fluorescence independent of the control system. To further investigate this effect, we analyzed the levels of Venus and mScarlet fluorescence independently in both the EcR-Cas9 reporter lines and no gRNA control populations. At 14 DPI, MF-treated plants of Line 3 exhibited a 44.3% decrease in Venus signal (*p* = 2.09 x 10^-7^) relative to the untreated plants (**Figure S10H**). A reduction in mScarlet fluorescence was also observed (16.8% decrease), although at a lower magnitude (**Figure S10I**). Although absolute fluorescence decreased for both fluorophores, the relative changes are consistent with net reporter activation by the control system regardless of the MF-associated effects on fluorescent output. Similar trends in Venus and mScarlet fluorescence were observed in other Cas9-EcR independent lines (**Figure S10B, S10C, S10E, S10F**). Furthermore, the fold changes in fluorescence, calculated as the ratio of induced fluorescence to the mean uninduced fluorescence, did not differ significantly between Venus and mScarlet in the no gRNA control population (**Figure S10D**). This finding further supports the interpretation that the relative fluorescence changes observed in the reporter lines reflect net activation by the control system. Collectively, these results demonstrate that the Cas9-EcR control system can achieve inducible activation in stable transgenic lines at levels comparable to transient assays

Given the MF-associated effects on fluorescent output, we next investigated whether lower MF concentrations could mitigate these effects while maintaining inducible activation (**Figure 4D**). We repeated the previous experiment in Line 3 using 0, 0.5, or 2.2 mM MF. Both induced conditions produced an increase in normalized signal, with the 2.2mM dose yielding a larger increase in fold change relative to the uninduced samples (2.78-fold increase, *p* = 2.0 x 10^-4^) and the 0.5mM dose (1.6-fold increase, *p =* 0.314; **Figure 4D**). We presume the smaller effect size of the 0.5mM dose led to a lack of significance when compared to the untreated samples. Both induced populations also displayed a reduced Venus signal relative to the uninduced samples (**Figure S10G**), with the 2.2mM treatment slightly lower than the 0.5mM treatment, although these differences were not statistically significant. Collectively, these results further support MF associated effects on fluorescent output, while also providing evidence consistent with dose-dependent responsiveness of the Cas9-EcR control system.

### Agrochemical-inducible CasG-based SynTFs transcriptionally reprogram endogenous and synthetic metabolic pathways

Thus far, we have demonstrated successful Cas9-EcR-based transgene activation. An additional utility of Cas9-based SynTFs is their ability to activate and repress both transgenes and endogenous loci (14, 28, 29), providing a powerful framework for reprogramming complex, multigene pathways that are difficult to engineer with conventional strategies. This capability could enable temporal control of metabolic genes to divert metabolic flux from endogenous to synthetic pathways. Such control is particularly important in cases where constitutive reprogramming of metabolism would impact fitness.

To test inducible metabolic reprogramming, we designed a genetic circuit to divert carbon flux, specifically caffeic acid, from the native lignin biosynthesis pathway into a synthetic pathway, the fungal bioluminescence pathway (FBP, **Figure 5B**, **5C**) (24, 38). In the FBP, hispidin synthase (Hisps) converts caffeic acid to Hispidin, which hispidin-3-hydroxylase (H3H) converts to luciferin. Luciferase (Luz) oxidizes luciferin, generating a high energy, light-emitting intermediate that caffeoyl pyruvate hydrolase (CPH) converts back to caffeic acid (**Figure 5C**) (27, 45). Because luminescence output depends on the coordinated activity of multiple enzymes, the FBP provides a useful platform for evaluating inducible control of multistep biosynthetic pathways.

**Figure 5.**
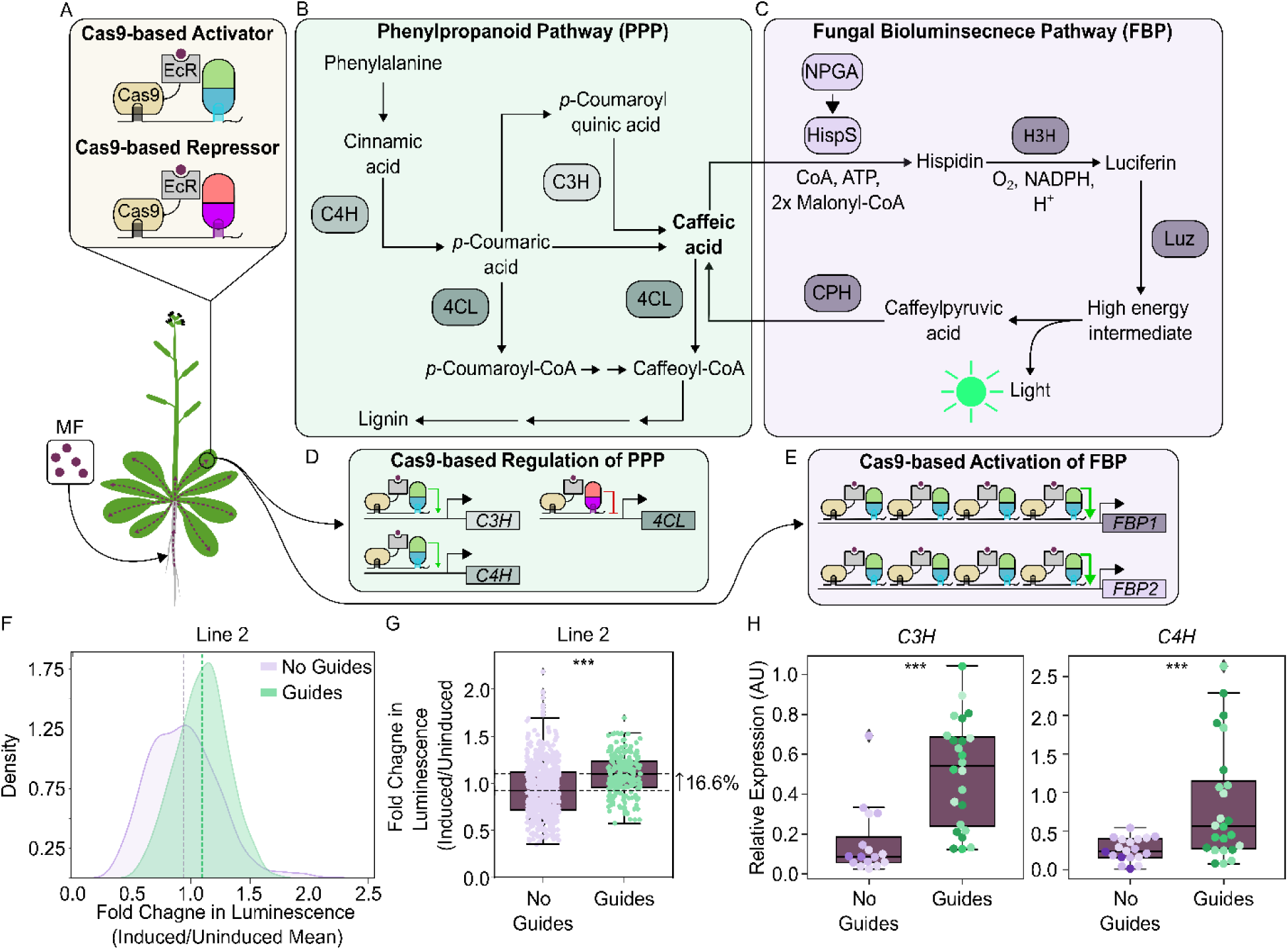
CasG-based, inducible redirection of metabolic flux. A) Schematic describing the Cas9-based control system. B, C) Schematic summarizing the key enzymatic steps of the phenylpropanoid pathway (PPP, B) and the Fungal Bioluminscence Pathway (FBP, C). D) Schematic representing the Cas9-based SynTF regulation implemented on the PPP to transcriptionally redirect metabolic flux into the FBP. E) Schematic representing the Cas9-based SynTF regulation implemented to activate expression of the FBP. F, G) Kernel density plots (F) and boxplots (G) representing the fold change in luminscence (induced sample/uninduced population mean) of plant populations with (n=215 measurements from Line 2) and without (n=578 measurements across 3 independent lines) PPP gRNAs. Each data point represents the fold change in luminsence (induced sample/uninduced population mean) of a single leaf. H) Boxplots representing the relative expression of *C3H* and *C4H* in leaf tissue of plants with (n=4 to 8 plants each from 4 independent lines) and without (n=3 to 8 plants each from 3 independent lines) PPP gRNAs and treated with MF. Each data point represents the relative expression from a sample of pooled leaf discs from a single plant with the color corresponding to plants from the same independent line. Asterisks represent results from a Welch’s two sample *t*-test (*p* < 0.05), * corresponds to *p* < 0.05, ** corresponds to *p* < 0.005, and *** corresponds to *p* < 0.0005.

To elucidate the ability of our system to regulate a multistep enzymatic pathway, we generated *A. thaliana* lines expressing the Cas9-EcR SynTF and the FBP under the control of 4xLacO pMin35s synthetic promoter (**Figure 5A**, **5E**). Luminescence measurements performed at 14 DPI revealed little to no difference between MF-treated plants relative to untreated controls. Plants were subsequently treated for an additional seven days, and luminescence measurements were repeated. Relative to un-treated controls, two of the three independent lines exhibited a modest increase in luminescence, although this increase was not statistically significant (**Figure S11**). Interestingly, one line showed a reduction in luminescence output upon treatment of MF, consistent with reductions observed in the fluorescent reporter system (**Figure S10, S11**). We reasoned that the lack of activation may be due to limited substrate availability as caffeic acid serves as a shared intermediate between the phenylpropanoid biosynthesis pathway (46) and the FBP (**Figure 5B**, **5C**), which may restrict pathway output. Therefore, we sought to enhance luminescent output by reprogramming expression of genes in the phenylpropanoid biosynthesis pathway to increase the available pools of caffeic acid.

Prior transient studies have shown that overexpression of caffeic acid biosynthesis genes and knockdown of enzymes that convert it into downstream products can alter metabolic flux to enhance the available pool of caffeic acid and hence increase FBP output (47). We implemented an inducible version of this metabolic reprogramming by generating *A. thaliana* lines expressing the FBP reporter system together with gRNAs designed to recruit the MCP-DREB2A activator to the promoters of *CYPS8A3* (*AtC3H*) and *CINNAMATE 4-HYDROXYLASE* (*AtC4H*) and the PCP-TPLN300 repressor to the promoter of *4-COUMARATE:COA LIGASE* (*At4CL*) (**Figure 5A**, **5D**). At 21 DPI, three independent lines containing the phenylpropanoid-targeted gRNAs (guides) displayed significantly increased luminescence relative to the untreated controls (**Figure S12A**). All lines displayed a distribution shift compared to the population without the phenylpropanoid-targeting gRNAs (No guides) when comparing the fold change in luminescence, calculated as the ratio of induced luminescence to the mean uninduced luminescence (**Figure 5F**, **S12B**). Line 2 showed the most significant shift, which represented a 16.6% increase in luminescence output when compared to the population without phenylpropanoid gRNAs (*p* = 1.35 x 10^-15^, **Figure 5G**). These results imply that our control system’s modulation of the phenylpropanoid pathway gene expression enables increased carbon flux to the FBP.

To confirm the increase in luminescence was associated with the modulation of phenylpropanoid pathway gene expression, we quantified transcript levels in treated and untreated plants from both the FBP-only and metabolic flux redirection lines. Across all lines tested, MF treatment resulted in a general decrease in absolute expression levels of *C3H*, consistent with the previously observed reduction in reporter fluorescence under induced conditions (**Figure S10, S13A**). However, baseline expression levels of *CH3* in untreated plants were comparable across genotypes and independent lines (**Figure S13B. S13D**), which supports valid cross-line and cross-genotype comparisons. To account for the MF-associated shifts in baseline expression, we therefore compared the relative expression for each line with the phenylpropanoid-targeting gRNAs to controls lacking them all under MF treatment. Transgenic lines carrying the phenylpropanoid-targeting gRNAs displayed a 6.2-fold and 2.3-fold increase in *C3H* and *C4H* relative expression, respectively, when compared to the population of transgenic lines with FBP-only gRNAs (**Figure 5H**). In contrast, no consistent decrease in *4CL* expression was observed in the targeted population (**Figure S13C**). Prior studies have shown this may be due to low gRNA efficiency (35). Together, these results demonstrate the utility of the Cas9-EcR system for inducible control of gene expression in multienzyme pathways spanning both native genes and transgenes to alter metabolic output.

### Agrochemical-induced CasG-based SynTFs can reprogram developmental signaling while mitigating pleiotropic effects

To further demonstrate the utility of the Cas9-EcR system, we applied it to temporal control of development. SynTF-mediated reprogramming of developmental pathways can engineer complex agronomic traits such as organ size (28–30). However, constitutive reprogramming may cause pleiotropic effects due to continuous pathway perturbation throughout development. We hypothesized that Cas9-EcR-based temporal control of expression could restrict pathway perturbation to selected developmental stages and mitigate these effects.

Previous work demonstrated that targeted activation of *DELLA* genes, which encode transcriptional regulators of the gibberellin (GA) signaling pathway can produce dwarfing (28). Such alterations in organ size were the basis of the green revolution, and hence control systems that these changes are agriculturally valuable. In this work, a nuclear-localized Cas9-based SynTF was guided to the promoters of the five *DELLA* genes to implement transcriptional activation. However, the potential developmental pleiotropies associated with constitutive *DELLA* activation, including impaired seed germination, were not investigated. Given the importance of GA signaling during germination (48, 49) and subsequent plant development, we sought to determine whether Cas9-based constitutive activation of *DELLAs* impairs germination and if temporal control using our Cas9-EcR-based control system could preserve germination while enabling transcriptional reprogramming at later developmental stages.

To test this, we adapted the previously established *DELLA* activation strategy by replacing the constitutively activate nuclear-localized Cas9 with our inducible Cas9–EcR system, enabling ligand-dependent control of SynTF activity (**Figure 6A**). We generated transgenic lines expressing Cas9–EcR with or without gRNAs targeting the five *DELLA* genes (**Figure 6A**) (28). Seeds from multiple constitutive and Cas9-EcR *DELLA* activation lines were sown on MS plates and germination was quantified at 2, 3, and 5 days post stratification. Across all timepoints, the Cas9-EcR *DELLA* activation lines exhibited significantly higher germination percentages than the constitutive activation lines, with a final germination that was 52.5% higher in the Cas9-EcR lines (*p* = 0.00013, **Figure 6B**). These results demonstrate that temporal control of SynTF activity can mitigate adverse effects of constitutive developmental pathway reprogramming on seed germination, supporting the utility of our inducible Cas9-EcR control system.

**Figure 6.**
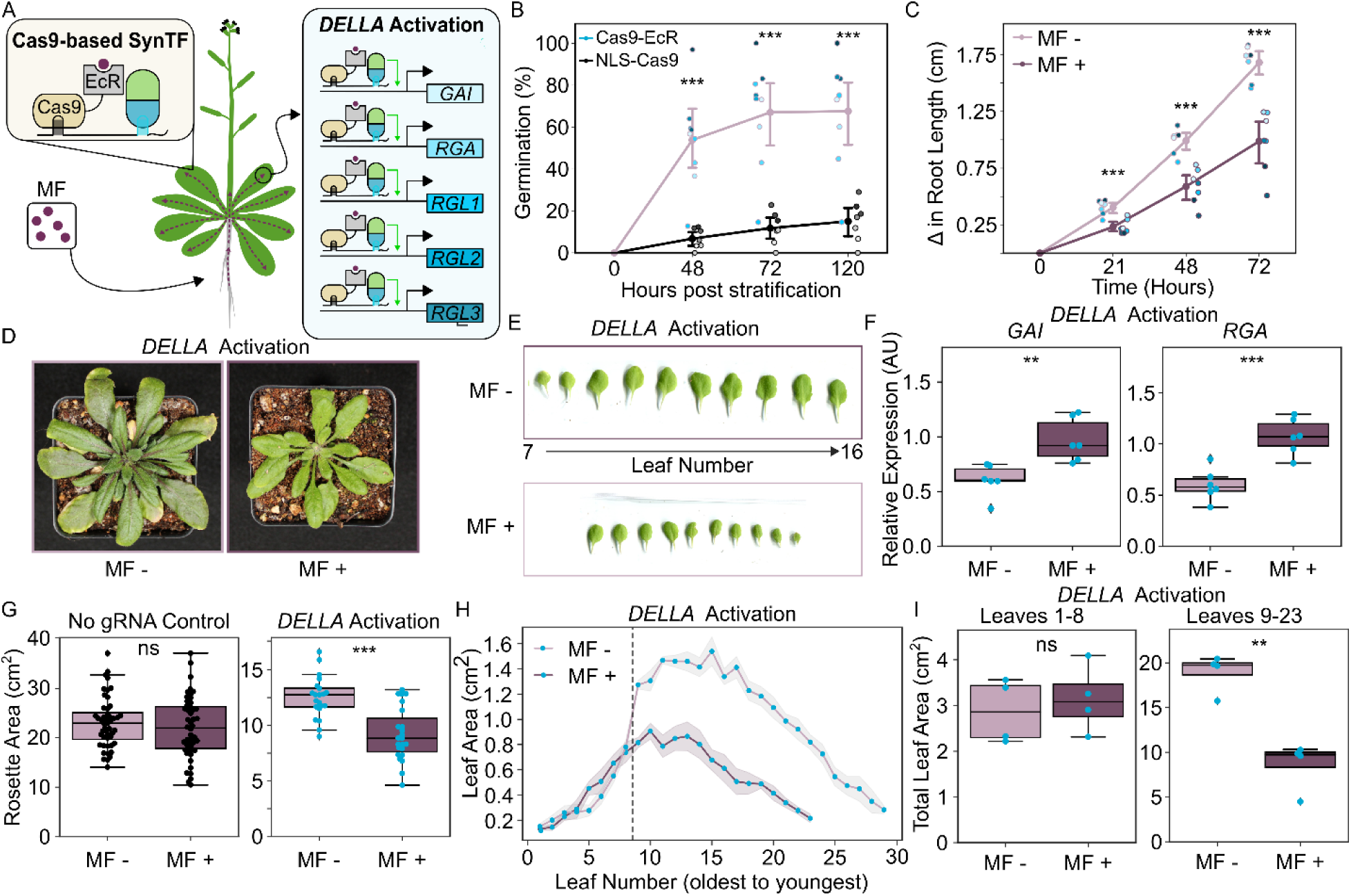
CasG-based, inducible control of plant development. A) Schematic summarizing the inducible Cas9-based control system for activation of five *DELLA* genes in *A. thaliana*. B) Line plots representing the percentage of seeds germinated from independent lines of the inducible (Cas9-EcR, n=9 indepednent lines) or constitutive (NLS-Cas9, n=7 independent lines) *DELLA* activation lines. Each dot represents the mean percentage of 3 replicates from a single independent line. C) Line plots representing the change in primary root length of Cas9-EcR *DELLA* activation lines (n=7 independent lines) treated with (dark magenta, MF+) or without (light magenta, MF-) MF for 72 hours. Each dot represents the mean change in root length of 6 to10 plants from a single independent line. D, E) Representative rosette (D) and leaf (E) images of *DELLA* activation plants treated with (dark magenta) or without (light magenta) MF. F) Boxplots representing the relative expression of *GAI* and *RGA* in *DELLA* activation Line 1 teated with (dark magenta, n=6 plants) or without (light magenta, n=6 plants) MF. G) Boxplots representing the rosette areas of a population of no gRNA control lines (left, n=2 independent lines, 24 plants per line per treatment) and a *DELLA* activation line (right, Line 1, n=21 to 23 plants per treament) treated with (dark magenta) or without (light magenta) MF. Each dot represents the rosette area of a single plant. H) Line plots representing leaf area across developmental age for plants from *DELLA* activation Line 1 treated with (dark magenta) or without (light magenta) MF. Each dot represents the mean leaf area across 4 plants. I) Boxplots depicting the total leaf are pre-(left, leaves 1-8, n=4 plants) and post-(right, leaves 9-23, n=4 plants) divergence from *DELLA* activation plants treated with (dark magenta) or without (light magenta) MF. Each dot represents a single plant. Asterisks represent results from a Welch’s two sample *t*-test (*p* < 0.05), * corresponds to *p* < 0.05, ** corresponds to *p* < 0.005, and *** corresponds to *p* < 0.0005.

To further evaluate the ability of our control system to temporally reprogram plant development, we investigated whether inducible gene regulation could enable dynamic control of organ size. We assessed seven independent transgenic lines by measuring primary root length after 72 hours of growth on MS agar plates with or without MF. All lines tested resulted in significantly reduced root growth in the presence of MF relative to untreated seedlings, a response that was not observed in the control population (**Figure 6C, Figure S14A**). The magnitude of root growth inhibition varied among independent lines, with reductions ranging from 26.7% to 72.3% compared to corresponding untreated controls (**Figure 6C**). This variation may reflect differences in T-DNA insertion or transgene expression among lines.

To further characterize inducible control of organ size, we conducted a soil-based induction assay using two independent lines that displayed differing degrees of responsiveness in the plate assay (Line 1 – 32.1% reduction in root length, Line 2 – 72% reduction in root length). MF treatment resulted in a 27% (*p* = 6.55 x 10^-6^) and 31% (*p* = 0.018) reduction in mean rosette size in the *DELLA* activation Lines 1 and 2, respectively (**Figure 6D**, **6G, S14G, S14H**). No significant reduction in rosette size was observed in the no gRNA control lines (*p* = 0.87, **Figure 6G, S14B**). Additional phenotypic characterization revealed reductions in both the size and the number of expanded leaves of MF-treated *DELLA* activation plants and displayed a distinct developmental timepoint at which growth began to diverge from untreated plants (**Figure 6E**, **6H, S14F**). Neither of these patterns were observed in no gRNA control lines (**Figure S14C, S14D**). To quantify overall growth across plants of each genotype, we calculated the total leaf area across leaves pre-(leaves 1 through 8) and post-(leaves 9 through 23) growth divergence that were harvested at a single timepoint (**Figure 6E, S14E**). Pre growth divergence, no differences in total leaf area were observed between the MF-treated and untreated plants for either the *DELLA* activation lines or the no gRNA control lines (**Figure 6I, S14E**). Post divergence, MF-treated *DELLA* activation lines displayed a significant reduction in total leaf area (54.7% reduction) relative to the untreated plants (*p* = 0.0012, **Figure 6I**). There was no reduction in cumulative growth observed in the no gRNA control lines post divergence (**Figure S14E**). *DELLA* activation lines treated with MF also had a reduction in the number of emerged and expanded leaves relative to the untreated plants, another result not observed in the no gRNA control lines (**Figure S14F**). These observations are consistent with previous reports demonstrating that elevated DELLA accumulation suppresses cell expansion and plant growth (50–52).

To further validate that the observed dwarfing phenotypes were associated with modulation of target gene expression, we quantified transcript levels of the two major *DELLA* genes, *GAI* and *RGA*, in leaves collected from *DELLA* activation Line 1. Treated plants exhibited increased *GAI* (1.60-fold increase, *p* = 6.74 x 10^-4^) and *RGA* (1.79-fold increase, *p* = 5.5 x 10^-3^) expression relative to untreated plants (**Figure 6F**). Neither *GAI* nor *RGA* expression levels were significantly elevated in the treated no gRNA control plants relative to untreated plants (**Figure S14I**). Additionally, MF treatment had no effect on gene expression levels of our housekeeping normalization control, PP2A (**Figure S14J**). Interestingly, the untreated *DELLA* activation plants from Line 1 were smaller than the untreated control plants (**Fig. S14G, S14B**), which is likely due to random T-DNA integration affecting growth in these lines independent of the GA signaling pathway, as the absolute levels of *GAI* and *RGA* were not significantly elevated in the untreated *DELLA* activation lines compared to the untreated no gRNA control lines (**Figure 6F, S14I**). Together, these results provide evidence that MF induces Cas9-EcR-based transcriptional reprogramming of a hormone pathway, resulting in activation of *DELLA* genes and the associated dwarfing phenotypes.

## Discussion

In this work, we establish a framework for designing integrated and programmable agrochemical-inducible expression control systems, characterize their performance *in planta*, and demonstrate their application for chemical control of metabolism and development. Our work demonstrates that the re-engineered ecdysone receptor (EcR) can be integrated with both Cas9-based SynTFs and site-specific DNA recombinases. This highlights the modularity of this signaling domain and extends the use of MF-responsive systems for both dynamic, dose-dependent and binary, switch-like induction. Our work builds on prior studies that used simpler control systems to achieve systemic induction of single genes (4, 53, 54) and demonstrates how more sophisticated control systems that respond to MF enable the dynamic regulation of multigenic metabolic and developmental pathways.

We selected EcR because prior work demonstrated selective responsiveness to MF formulated as Intrepid2F for field stability and systemic vasculature distribution (4, 5, 53, 54). This has been demonstrated via root drench delivery (4, 53, 54), which overcomes limitations of foliar application where the cuticle can restrict inducer uptake and reduce induction uniformity (55, 56). Our demonstration of reproducible induction of gene expression in systemic tissues following root-drench application, across both transient assays and stable transgenic lines, is consistent with previous work and validates the utility of MF as a whole-plant inducer.

We also observed that MF, at the concentrations used, impacts global gene expression independent of the control system. This effect was not reported in a previous study using Intrepid2F at concentrations approximately 30-fold higher than those used here (57), although its induction durations were substantially shorter. This difference suggests that exposure duration, rather than concentration alone, may determine this effect. Despite this, plants showed no MF-associated growth-defects, suggesting that the inducer’s effects are context and trait dependent and may not substantially impact other developmental phenotypes, such as yield. Field studies will be critical to identify induction regimes relevant to agricultural applications, given the differences between our concentrations and application modalities and those used by growers.

Our initial characterization revealed the presence of both incomplete activation and basal leak when EcR was integrated with both the Cas9-based and the recombinase-based control systems. Incomplete activation may be due to either lower functional expression of the control system when compared to Cas9-NLS or submaximal inducer concentrations in the sampled tissue. The basal leak observed is consistent with previous reports of leakiness in ligand-gated control systems (5, 23, 58, 59). To reduce leak and improve the dynamic range, we engineered the nuclear trafficking dynamics of each system through the incorporation of an NES. This strategy reduced leaky activation in the uninduced state in both systems. The resulting constructs also enabled direct comparison of NES strength. Each NES decreased basal leak at varying degrees, suggesting that modulation of nuclear export could serve as a useful strategy for tuning gene regulation in future control systems. More broadly, this assay provides a framework for quantitatively comparing the relative strengths of additional NES sequences *in planta*.

While similarly reducing basal leak, NES incorporation had markedly different effects on induced activity in the two systems. Induction was completely abolished in the Cas9-based system, whereas maximal induction was only modestly reduced in the recombinase-based system. We attribute this difference to the distinct mechanisms by which each system transduces the MF signal. The recombinase-based system converts MF perception into an irreversible binary switch from an off-state, in which transgene expression is not possible, to an on-state, in which a strong constitutive promoter drives expression. This irreversibility makes the control system sensitive to a leaky off state because even rare nuclear import in the absence of inducer would accumulate over time. However, binary switching creates threshold-like behavior, where the strength of activation in the on-state is independent to the degree of nuclear import. As a result, incorporating the NES into the recombinase-based control system mitigated leak without strongly impacting the strength of activation in the on state. In contrast, the Cas9-based control system converts MF perception into a reversible and dose-dependent modulation of transcription. Because regulation depends on continuous presence of the SynTF in the nucleus, infrequent nuclear import in the absence of the inducer produces only limited regulation and does not accumulate overtime. However, this dose-dependency also means that a strong on-state necessitates maximum nuclear import in the presence of the inducer. This explains why engineering the basal rate of nuclear trafficking is a good strategy to improve the transduction properties for recombinase-based control systems, and why this strategy creates a tradeoff between leak and inducibility in Cas9-based control systems. This highlights the need for developing alternative engineering strategies that decouple this trade-off to improve the transduction properties of this control system in the future.

Although NES engineering improved the dynamic range in the recombinase-based control system, stable lines were susceptible to silencing over time. This is consistent with prior reports of hypermethylation at Cre-targeted loxP sites in plants (38). The recombinase-based control system is also limited to the regulation of transgenes, whereas Cas9-based systems can regulate both transgenes and native loci.

We demonstrated the utility of the inducible Cas9-based control system for dynamic control of multienzyme pathways by leveraging its flexibility to simultaneously activate the expression of FBP transgenes and native genes that drive substrate biosynthesis. These results provide a proof-of-concept that this control system can be applied to dynamically reprogram metabolic flux in plants. We further demonstrated the utility of being able to target native genes by using the Cas9-based system to dynamically modulate plant development via inducible regulation of genes in the GA signaling pathway. These results demonstrate that ligand-dependent SynTF systems can enable inducible control of complex developmental phenotypes, such as organ size, while also mitigating pleiotropies associated with constitutive perturbation. Taken together, our findings demonstrate that agrochemical-inducible controls systems provide a powerful and modular framework for the temporal control of both endogenous and synthetic pathways.

Future work should focus on improving the dynamic range of both the Cas9-based and recombinase-based control systems. This may include directed evolution of the EcR domain to decrease leak and the incorporation of stronger transcriptional effector domains such as MoonTag (60) or VPR (61) to enhance regulation strength of SynTFs. Regulation strength could also be improved through identification of more efficient gRNAs via rapid screening *in planta*, as has been demonstrated previously (29, 35). Furthermore, replacing the Cre recombinase with alternative site-specific recombinases, such as serine integrases (62), could reduce susceptibility of epigenetic transgene silencing. Additionally, integrating the two approaches into a single control system by using a recombinase to turn on expression of Cas9 could reduce leak while retaining the programmability of Cas9-based systems. Finally, achieving precision design of complex traits will require both temporal and spatial control of expression. Future applications of these tools could incorporate tissue specific promoters or ribozyme-based strategies to enable spatial restriction of activity (33). Ultimately, the ability to precisely control gene expression in both space and time will create new avenues to engineer complex traits and circumvent negative pleiotropies.

## Acknowledgments

We would like to thank Rich Conant for support from the Grantham Foundation, and Sinead Cahill from Devin O’Connor’s Lab at Colorado State University for assistance with confocal microscopy.

## Author Contributions

T.B. and A.K. planned and designed the research. T.B., L.F., S.S, J.V.B., and L.C. performed experiments. T.B. and L.F. analyzed the data, and T.B. and A.K. wrote the manuscript.

## Data Availability

All data analysis was performed in python using Jupyter Lab (version 3.4.4). All the raw data and code used to analyze the data is available on the following github repository (https://github.com/arjunkhakhar/260729_Methoxyfenozide_Control_Systems). Other data are included in the article and/or Supplemental Information.

