## Supplemental Information for "Agrochemical-responsive gene expression control systems for modulating plant development and metabolism"

### Supplemental Materials and Methods

#### Plasmid construction

All plasmids constructed in this work were built using Golden Gate Assembly and Modular cloning (24–26) and are listed with links to their annotated plasmid maps in **Supplemental Table S1**. Most DNA parts (promoters, CDS, terminators) were amplified from previously reported plasmids with primers that added flanking restriction sites and compatible overhangs (24, 27–31). The $CfEcR_{VY}$ domain was synthesized (Twist Bioscience) with flanking BsaI sites and compatible overhangs for assembly. Guide RNA (gRNA) target sites were designed using CHOPCHOP v3 (32) and the target sequences are listed in **Supplemental Table S2**. The gRNA and nuclear export sequences were added to each DNA part using primers and stitched together during the assembly and ligation process. All plasmids used in this work will be available via AddGene.

#### Transient assays

Cultures of *Agrobacterium tumefaciens* strain GV3101 carrying the SynTF and/or reporter plasmids were inoculated into 5mL of LB supplemented with Gentamycin (30mg/L) and Kanamycin (50mg/L). Cultures were grown for ~20 hours at $28^0$C and 250 rpm. Cultures were centrifuged for 10 minutes at 3000xg, resuspended in infiltration medium (10mM MES, 10mM MgCl2, pH 5.6) to an OD600 of 0.6. After dilution, acetosyringone was added to each culture at a final concentration of 0.2 mM and incubated at room temperature for 2 to 4 hours. Samples were infiltrated into the abaxial side of *Nicotiana benthamiana* leaves using a 1 mL needless syringe. For Cas9-EcR transient assays, immediately prior to infiltration, co-cultures were made by mixing a 1:1 ratio of the SynTF plasmid and the reporter plasmid. For EcR-Cre transient assays, co-cultures were not necessary as all control system components were co-expressed on the same T-DNA. Infiltrated plants were returned to a Percival growth chamber programmed to the following conditions: $25^0$C, 12-hour light/12-hour dark, 70% relative humidity. Root drenches were conducted by top watering each plant with 500 mL of MF or tap water. MF was prepared by mixing 3.4 mL of Intrepid 2F into 996.6 mL of deionized water.

**Stable line generation**

All stable *Arabidopsis thaliana* transgenic lines in this work were generated using floral dip (10). For the Cas9-EcR control system, the Cas9-EcR T-DNA plasmids were first transformed into *A. thaliana*. T0 seeds were collected, sterilized with 70% ethanol for 20 minutes, washed once with 95% ethanol and sown on 1/2x MS agar plates containing Kanamycin (50mg/mL). Seeds were stratified on plates at $4^{0}$C in the dark for 72 hours then place into a Percival growth chamber. Positively selected seedlings were transplanted into soil and grown to produce T1 seed. This process was repeated to generate T2 seeds. Cas9 expression was quantified in T2 lines using RT-qPCR to identify a high expressing Cas9 line. T2 seeds of a high expressing Cas9 line were germinated and transplanted into soil as described above. The ratiometric fluorescent reporter T-DNA with and without the gRNAs targeting mScarlet for activation were transformed into multiple T2 Cas9-EcR plants. For the FBP lines, a T-DNA vector that encoded the Luz-H3H-CPH (FBP1) fusion protein driven by the 4xLacO pMin35s promoter, the Hisps-NPGA (FBP2) fusion protein driven by the 4xLacO pMin35s promoter, a gRNA-MS2 scaffold targeting the LacO elements upstream of the pMin35s promoter, and a module with or without gRNAs targeting the phenylpropanoid pathway were transformed into multiple T2 Cas9-EcR plants. T0 seeds were sterilized as described above and germinated on dual selection 1/2x MS agar plates containing Kanamycin (50mg/mL) and Glufosinate-ammonium (15mg/mL). Independent lines were taken to the T1 generation and used for further experiments. Cre recombinase T-DNA plasmids with and without the reporter were transformed into *A. thaliana* and taken to seed. T0 seeds were sterilized and germinated on 1/2x MS agar plates containing Kanamycin (50mg/mL) as described above. T1 seeds were collected and used for further experiments.

**Methoxyfenozide induction of stable transgenic lines**

Seeds of EcR-Cre and EcR-Cre HIV1 REV independent transgenic lines were sterilized and plated as described above. Plates were stratified in the dark at $4^{0}$C for 3 days. Plates were subsequently placed vertically in a Percival growth chamber set to short day conditions

(9/15-hour light/dark cycle) and 22$^{0}$C. Up to 10, 10-day old seedlings were transferred to 1/2x MS agar plates supplemented with 80nM methoxyfenozide (Milipore Sigma, Product Number 32507) or DMSO as a negative control. Fluorescence was quantified at multiple timepoints as described below. Root length was quantified at 0, 21, 48, and 72-hour timepoints by scanning the plates on a flatbed scanner (Epson). Total root length for each timepoint was measured in Fiji and the difference in primary root length for T21, T48, and T72 were calculated by subtracting the initial primary root length at T0.

Seeds for all the *A. thaliana* lines containing Cas9-EcR were sterilized with 70% ethanol for 20 minutes, washed once with 95% ethanol, and sown on 1/2x MS containing 0.8% PhytoAgar™ (plantmedia Cat. No. 401000721) and supplemented with Kanamycin (50mg/L) and Glufosinate-ammonium (15mg/L). Seeds were stratified on plates at 4$^{0}$C in the dark for 72 hours and subsequently place into a Percival growth chamber set to short day conditions (9/15-hour light/dark cycle) and 22$^{0}$C. Cre recombinase and *DELLA* activation lines were sterilized and germinated on 1/2x MS agar plates and supplemented with Kanamycin (50 mg/L) as described above. For the demethylation experiment, seeds of EcR-Cre and EcR-Cre HIV1 REV lines were germinated on 1/2x MS agar plates supplemented with Kanamycin and 50uM 5-Azacitidine (AzaC). Plates were left in the chamber for 14 days, transplanted into soil (1:1 ProMix HP/Greensgrade) in 2.5-inch square pots, and placed in a Conviron growth chamber set to short day conditions (9-hour light/15-hour dark) at 22$^{0}$C. Plants were established in soil for two weeks and were bottom watered three days per week. After two weeks, a subset of the plants was top watered twice per week with either 31 mL of 2.2 mM methoxyfenozide as Intrepid 2F or tap water. After 14 days of induction (4 total applications), fluorescence, luminescence, or shoot phenotypes were quantified.

**Quantification of reporter fluorescence**

For all transient assays, reporter fluorescence was quantified using a Tecan Spark plate reader. Leaf discs were harvested from infiltration sites of *N. benthamiana* transient assays or rosette leaves from *A. thaliana* stable lines and placed in the bottom of 96 well plates (Corning, black polystyrene, clear bottom) adaxial side up. Each well contained 10 uL of water to adhere leaf discs to the bottom of each well. Venus and mScarlet measurements were performed using the following settings: Venus excitation 505 nm, Venus emission 540 nm, Venus bandwidth 15 nm, mScarlet excitation 560 nm, mScarlet emission 605 nm, mScarlet bandwidth 20 nm. Both fluorescence measurements were read from the bottom to capture the signal on the abaxial side of the lead. For each experiment, the gain and z-position were first measured using the optimal setting and then manually set across all plates for that experiment.

For methoxyfenozide plate assays, mScarlet fluorescence was quantified using an Analytik Jena UVP ChemStudio PLUS Imaging System. Image acquisition settings were held constant across all samples, including capture binning (1), epi lighting (green), trans lighting (off), brightness (50%), emission filter (605 nm), and exposure time (15 seconds). Fluorescence intensity was quantified in Fiji (ImageJ2) by measuring the mean gray value within a fixed-size region of interest (ROI) for each seedling. To correct for image-to-image variation in background fluorescence, ten background ROIs were measured for each image, and their mean intensity was calculated. Background-corrected fluorescence was determined by subtracting the mean background intensity from the mean intensity of each seedling ROI.

**Quantification of luminescence output**

Reporter luminescence was quantified using a AnalytikJena UVP ChemStudio PLUS Imaging System and VisionWorks imaging software. Camera focus was manually adjusted for every set of plants. Image acquisition settings were held constant across all samples, including capture binning (4), epi lighting (off), trans lighting (off), and brightness (100%). Plants were imaged from the top view with 10 minutes of exposure. Resulting images were analyzed using the VisionWorks Area Density tool. Leaf area was outlined using the ROI drawing tool with each leaf as an individual ROI. To account for differences in images, four independent ROIs were measured for background mean density. Luminescence was quantified by subtracting the mean background density from the mean raw density.

***Arabidopsis* shoot and germination phenotyping**

Phenotyping of *A. thaliana* shoots was performed at 14 DPI. Individual plants were photographed using a Canon EOS Rebel T6 camera and rosette area estimates were measured in Fiji by tracing the outer perimeter of the rosette. Four plants from each independent line from the no gRNA control and *DELLA* activation lines were randomly selected for more in-depth shoot phenotyping. Each plant was dissected by leaf and scanned on a flatbed scanner (Epson) in order of developmental age. Individual leaf areas were measured in Fiji.

For evaluating germination efficiency of the *DELLA* activation lines, independent lines of the Cas9-EcR *DELLA* activation (n=9) and global *DELLA* activation lines (n=7) were sown on 1/2x MS agar plates as described above. The total number of seeds germinated was collected at 48, 72, and 120 hours after stratification. The germination efficiency of each genotype was measured by calculating the percent of seeds germinated at each timepoint for each independent line. Three biological replicates were completed for each independent line.

**Gene expression analysis**

The relative expression of target genes was measured using a modified version of a previously reported protocol (5, 11). For characterization of *C3H*, *C4H*, and *4CL*, leaf discs of MF-treated and untreated plants were collected and immediately frozen in liquid nitrogen. For characterization of *GAI* and *RGA* expression, whole leaves of the developmentally same age were harvested and immediately frozen in liquid nitrogen. RNA was extracted using the QIAGEN RNeasy Plant Mini Kit (Cat No. 74904). The RNA samples were treated with the TURBO DNA-*Free*™ kit (Invitrogen Cat No. AM1907) to remove genomic DNA. Complementary DNA (cDNA) was synthesized using the LunaScript® RT primer free MasterMix kit with gene specific primers (GAI and RGA) or the SuperMix Kit (*C3H*, *C4H*, and *4CL*). For *GAI* and *RGA*, proper removal of genomic DNA was confirmed with PCR using GoTaq Green Master Mix (Promega Cat No M7122) and primers that span the first and second intron of *GID1a* (5). For *C3H*, *C4H*, and *4CL*, proper removal of genomic DNA was confirmed using an RT negative control during the qPCR reaction. Concentrations of all genes were quantified using RT-qPCR performed using Luna® Universal qPCR Master Mix on a BioRad

CFX Opus qPCR machine. Cycle threshold values for all genes were normalized to the housekeeping gene, *PP2A*. Gene specific, PCR, and RT-qPCR primers are listed in **Supplemental Table S4**.

**Data analysis**

All data analysis was performed in Python using Jupyter Lab (version 3.4.4). The *p*-values reported were calculated using either an ANOVA followed by a Tukey HSD test or Welch's *t*-test for pairwise comparisons, where appropriate, using the scipy package in python. All the data that is presented was plotted using the seaborn package and functions from the matplotlib library (54). All the raw data and code used to analyze the data is available on the following github repository:
https://github.com/arjunkhakhar/260729_Methoxyfenozide_Control_Systems.

### Supplemental Tables

**Supplemental Table S1.** Plasmid names, components, and maps of plasmids used in this work.

| Plasmid Number | Plasmid Use | Plasmid Components | Plasmid Map |
|---|---|---|---|
| P282 | NLS-Cas9 SynTF Positive Control | pTRANS_220d-pUBQ10:NLS-AtCas9:tHSP-pGmUbi:NLS-MCP-DREB2A-tUBQ1-p35s:NLS-PCP-TPLN300:tNos | P282 |
| P299 | Cas9-EcR SynTF | pTRANS_220d-pUBQ10:SpCas9-CfEcR:tHSP-pGmUbi:NLS-MCP-DREB2A-tUBQ1-p35s:NLS-PCP-TPLN300:tNos | P299 |
| P843 | Cas9-EcR BR1 NES SynTF | pTRANS_220d-pUBQ10:EcR-AtCas9-BR1_NES:tHSP - pGmUbi:NLS-MCP-DREB2A-tUBQ1 - p35s:NLS-PCP-TPLN300:tNos | P843 |
| P844 | Cas9-EcR HIV1 REV NES SynTF | pTRANS_220d-pUBQ10:EcR-AtCas9-HIVREV1_NES:tHSP - pGmUbi:NLS-MCP-DREB2A-tUBQ1 - p35s:NLS-PCP-TPLN300:tNos | P844 |
| P288 | Ratiometric reporter with gRNAs | pTRANS_230d-pUBQ1mut:Venus:tHSP-4xLacO-pMinimal 35S:mScarlet:tHSP-pAtU6:LacO_targeting_truncated_gRNA_1xMS2:tU6 | P288 |
| P291 | Ratiometric reporter without gRNAs | pTRANS_230d-pUBQ1mut:Venus:tHSP-4xLacO-pMinimal 35S:mScarlet:tHSP | P291 |
| P849 | EcR-Cre | pTRANS_220d – pUBQ1mut:NLS-Venus-tHSP – tNOS_antisense-lox66-pUBQ10_antisense-lox77:mScarlet:tHSP-pGmUbi:EcR-Cre_intron:tUBQ1 | P849 |
| P847 | EcR-Cre BR1 NES | pTRANS_220d – pUBQ1mut:NLS-Venus-tHSP – tNOS_antisense-lox66-pUBQ10_antisense-lox77:mScarlet:tHSP - pGmUbi:EcR-Cre_intron-BR1_NES:tUBQ1 | P847 |
| P848 | EcR-Cre HIV1 REV NES | pTRANS_220d – pUBQ1mut:NLS-Venus-tHSP – tNOS_antisense-lox66-pUBQ10_antisense-lox77:mScarlet:tHSP - pGmUbi:EcR-Cre_intron-HIVREV_NES:tUBQ1 | P848 |
| P319 | Cre unactivated reporter | pTRANS_230d-pUBQ1mut:Venus:tHSP-tNOS_antisense-lox66-pUBQ10_antisense-lox77:mScarlet:tHSP | P319 |
| P1170 | Cas9-EcR FBP activation | pTRANS_210d-p35Smin_FBPF2_tHSP-p35Smin_FBPF1_tOCS-pAtU6:LacO_targeting_truncated_gRNA_1xMS2:tU6 | P1170 |

| Plasmid Number | Plasmid Use | Plasmid Components | Plasmid Map |
|---|---|---|---|
| P1173 | Cas9- EcR FBP activation + Phenylpropanoid gRNAs | pTRANS_210d-p35S_C3H-C4H-activation-2xMS2_4CL-repression-2xPP7_t35S-p35Smin_FBPF2_tHSP_p35Smin_FBPF1_tOCS-pAtU6:LacO_targeting_truncated_gRNA_1xMS2:tU6 | P1173 |
| P514 | Cas9-EcR No gRNA control | pTRANS_220d AtUBQ10:Cas9-EcR-tHSP-pGmUBI:NLS-MCP-DREB2A-p35s:PCP-TPL300-tNOS | P514 |
| P513 | Cas9-EcR DELLA activation | pTRANS_220d-AtUBQ10:Cas9-EcR-tHSP-pGmUBI:NLS-MCP-DREB2A-p35s:PCP-TPL300-tNOS-p35s:AtDELLA_guides1-1xMS2:t35s | P513 |

**Supplemental Table S2.** List of guide RNAs targets, uses, and target sequences used in this work.

| **Target** | **Use** | **gRNA target sequence (5'>3')** |
|---|---|---|
| LacO element | Activation of 4xLacO pMin35s synthetic promoter | CTAGAAAGAAGAAA |
| *C3H* promoter | PPP redirection into FBP | GGTCAGGTACATGT |
| *C4H* promoter | PPP redirection into FBP | AAGAGTGAGAACGA |
| *4CL* promoter | PPP redirection into FBP | AGGTTGATTTTATC |
| *GAI* promoter | *DELLA* activation | TGTCATGCAACTAG |
| *RGA* promoter | *DELLA* activation | CATCTAAAGTGATA |
| *RGL1* promoter | *DELLA* activation | TTGCGGTTGTACCA |
| *RGL2* promoter | *DELLA* activation | TTCTTAAACCCAAA |
| *RGL3* promoter | *DELLA* activation | CGGGCTGTTACTTT |

**Supplemental Table S3.** List of primer names, gene targets, and sequences used for PCR or qPCR.

| Primer name | Gene target (Accession Number) | Primer sequence (5'>3') |
|---|---|---|
| qPCR-[C3H 1]-f | C3H (AT2G40890) | GATGGACACGACAGCGATAA |
| qPCR-[C3H 1]-r | | GAGAAATCTGCCTCGGTTAAGA |
| qPCR-[C4H 1]-f | C4H (AT2G30490) | AGCTACCTCCAGGTCCTATAC |
| qPCR-[C4H 1]-r | | CGCCGAATTTCTTAGCGTAATC |
| qPCR-[4CL1 1]-f | 4CL (AT1G51680) | CGGAGACGGAGAAGTATGATTTG |
| qPCR-[4CL1 1]-r | | CCTGACCGAGTTTGGCATTA |
| qPCR-[AtGAI]-f | GAI (AT1G14920) | CTATGCTCACCGACCTTAATCC |
| qPCR-[AtGAI]-r | | GACGAAGAAGCCGAATCGATAG |
| qPCR-[AtRGA1]-f | RGA (AT2G01570) | CGCAGATTGGTGGAGTCATAG |
| qPCR-[AtRGA1]-r | | CTTGCGAGTCAACCAGGATAA |
| [mScarlet]-r | Orientation specific PCR | GATCCTTCCATGTGGACCTTAAA |
| [AtUBQ10]-f | | GCTCTTCTTCTTCACAATTCAGATTTC |

### Supplemental Figures

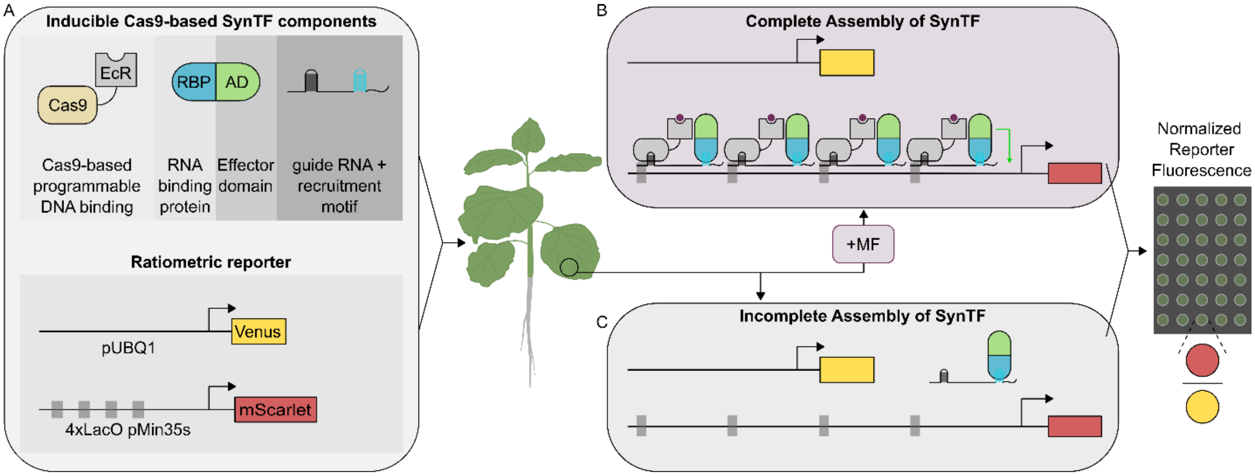


**Supplemental Figure S1. Transient agroinfiltration assay for the Cas9-based control system.** A) Schematic depicting the components of the two T-DNA plasmids co-infiltrated into *Nicotiana benthamiana*. B) Control system assembly in the presence of methoxyfenozide (MF). C) Lack of control system assembly in the absence of MF.

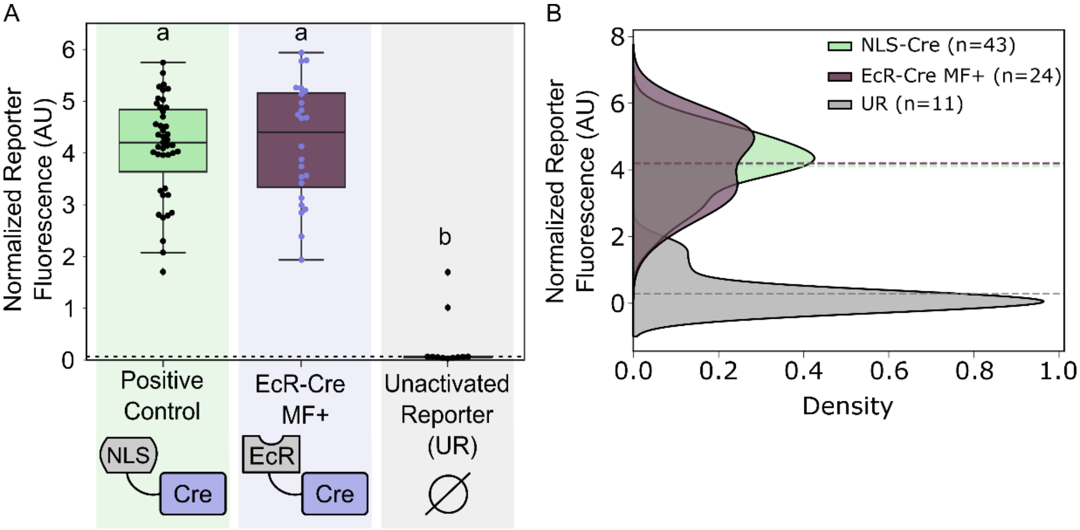


**Figure S2. Demonstration of complete activation of the EcR-Cre control system.** A) Boxplots representing the normalized reporter fluorescence from transient assays of plants infiltrated with the EcR-Cre control system and treated with (dark magenta, MF +, n=10 leaves across 3 independent plants) compared to a nuclear localized Cre (green, positive control) and an unactivated reporter (gray, n=20 leaves across 6 independent plants). B) KDE plots representing the distributions of the normalized reporter fluorescence for the EcR-Cre (dark magenta), NLS-Cre, and an unactivated reporter (UR). The vertical dashed lines represent the mean normalized reporter fluorescence for each version of the system. The total number of fluorescence measurements (leaf discs) is stated in the figure legend. Different letters represent statistically significant differences (One-way ANOVA followed by Tukey HSD test, $p < 0.05$).

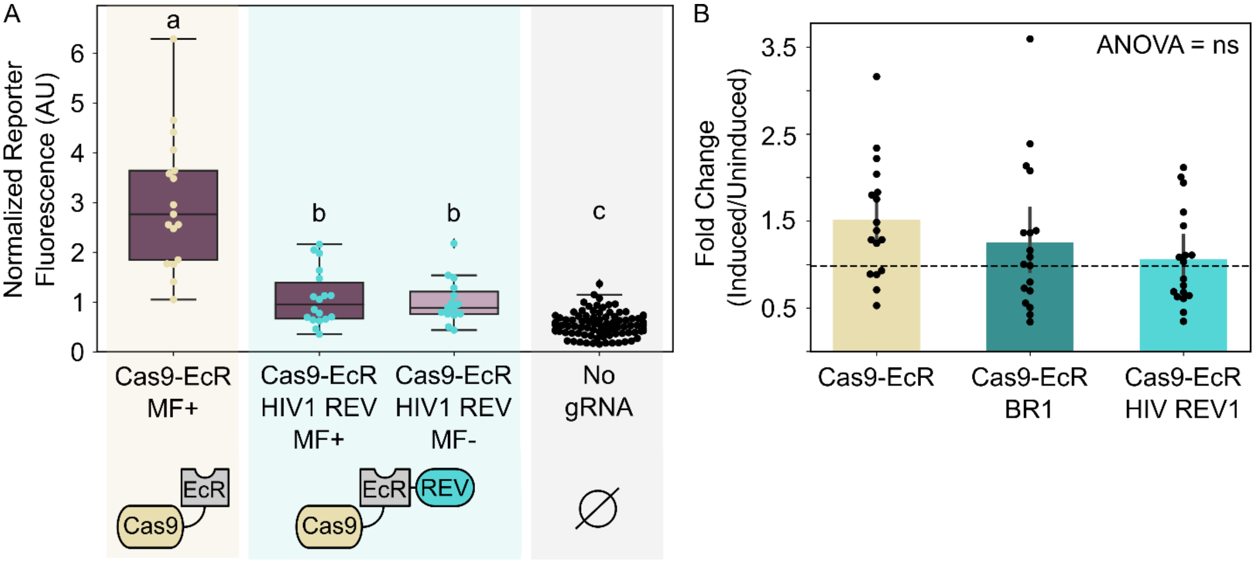


**Supplemental Figure S3. Engineering nuclear trafficking dynamics in the Cas9-based control system.** A) Boxplots representing the normalized reporter fluorescence of plants co-infiltrated with the Cas9-EcR or Cas9-EcR HIV1 REV control systems. Plants were either treated with (dark magenta, MF +, n=6 leaves across 3 independent plants) or without (light magenta, MF-, n=6 leaves across 3 independent plants) MF. B) Barplots representing the fold change in normalized reporter fluorescence of the induced samples to the uninduced samples for the Cas9-EcR (n=10 leaves across 3 independent plants), Cas9-EcR BR1 (n=10 leaves across 3 independent plants) and the Cas9-EcR HIV1 REV (n=10 leaves across 3 independent plants) control systems. Across all plots, every data point represents a normalized fluorescence measurement from a single leaf disc. Different letters represent statistically significant differences (One-way ANOVA followed by Tukey HSD test, $p < 0.05$).

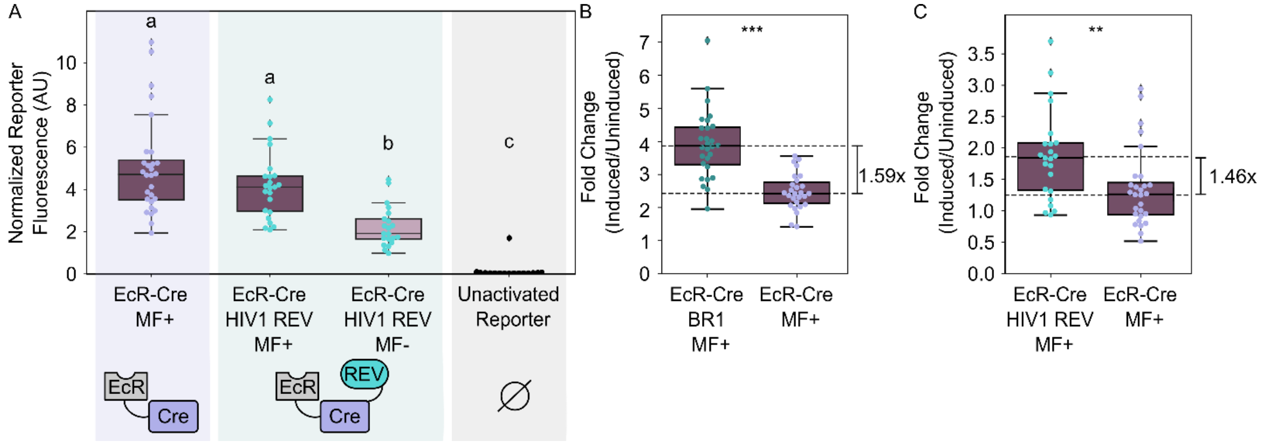


**Supplemental Figure S4. Engineering nuclear trafficking dynamics in the recombinase-based control system.** A) Boxplots representing the normalized reporter fluorescence of plants infiltrated with the EcR-Cre (purple) or EcR-Cre HIV1 REV (light teal) control systems. Plants were either treated with (dark magenta, MF +, n=10 leaves across 3 independent plants) or without (light magenta, MF-, n=10 leaves across 3 independent plants) MF. B) Boxplots representing the fold change in normalized reporter fluorescence (induced/uninduced mean) EcR-Cre BR1 (dark teal) and EcR-Cre (purple) control systems. C) Boxplots representing the fold change in normalized reporter fluorescence of the induced samples to the uninduced samples for the EcR-Cre HIV1 REV (light teal) and EcR-Cre (purple) control systems. Across all plots, every dot of the same color corresponds to an independent biological replicate. Different letters represent statistically significant differences (One-way ANOVA followed by Tukey HSD test, $p < 0.05$). Asterisks represent results from a Welch's two sample *t*-test ($p < 0.05$), * corresponds to $p < 0.05$, ** corresponds to $p < 0.005$, and *** corresponds to $p < 0.0005$.

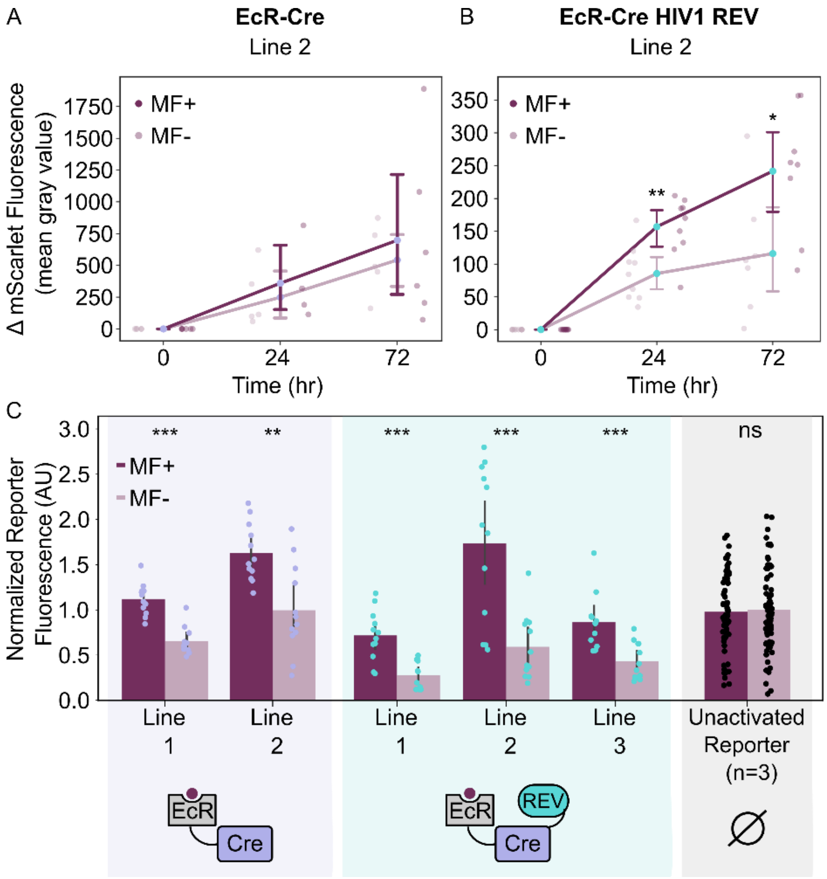


**Supplemental Figure S5. Characterization of the Cre-recombinase control system in stable transgenic lines.** A, B) Line plots representing the mScarlet fluorescence over 72 hours from an EcR-Cre (A, Line 2, n=6 plants per treatment) or EcR-Cre HIV1 REV (B, Line 2, n=10 plants per treatment) *A. thaliana* independent transgenic line treated with (dark magenta, MF +) or without (light magenta, MF -) MF. C) Barplots representing the normalized reporter fluorescence of the EcR-Cre (purple, n=4 plants per line per treatment), EcR-Cre HIV1 REV (light teal, n=4 plants per line per treatment), and the unactivated reporter lines (black, n= 3 independent lines, 3 plants per line per treatment) treated with (dark magenta) or without (light magenta) MF. Asterisks represent results from a Welch's two sample *t*-test ($p < 0.05$), * corresponds to $p < 0.05$, ** corresponds to $p < 0.005$, and *** corresponds to $p < 0.0005$.

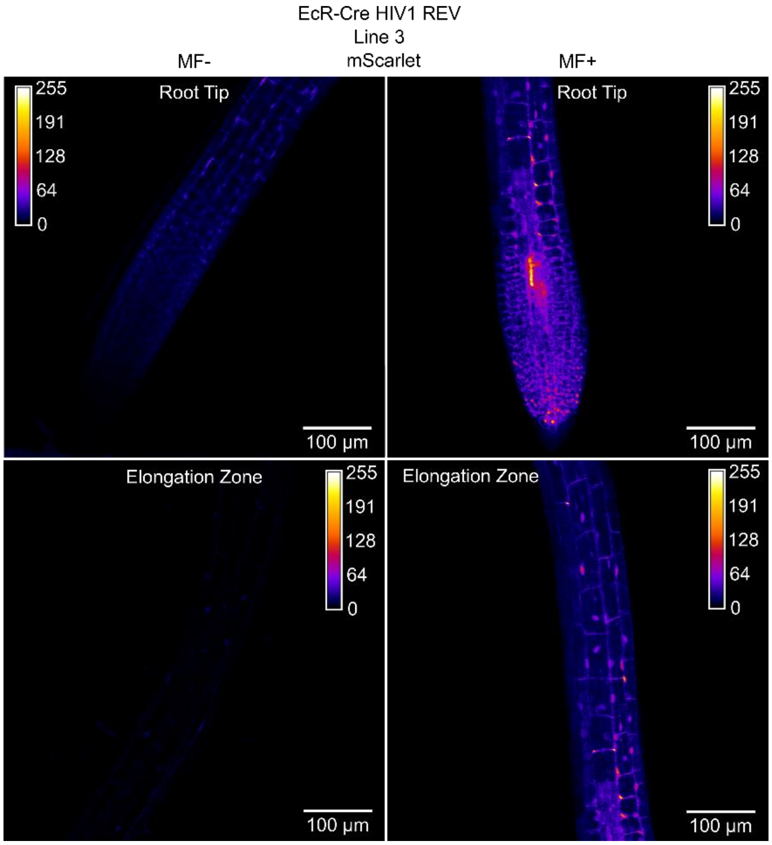


**Supplemental Figure S6. Characterization of the Cre-recombinase control system in stable transgenic lines.** Confocal images of the root tip (top) and elongation zone (bottom) of different biological replicates from an EcR-Cre HIV1 REV transgenic line (Line 3) that were treated with (right) or without (left) MF at 162 hours post induction.

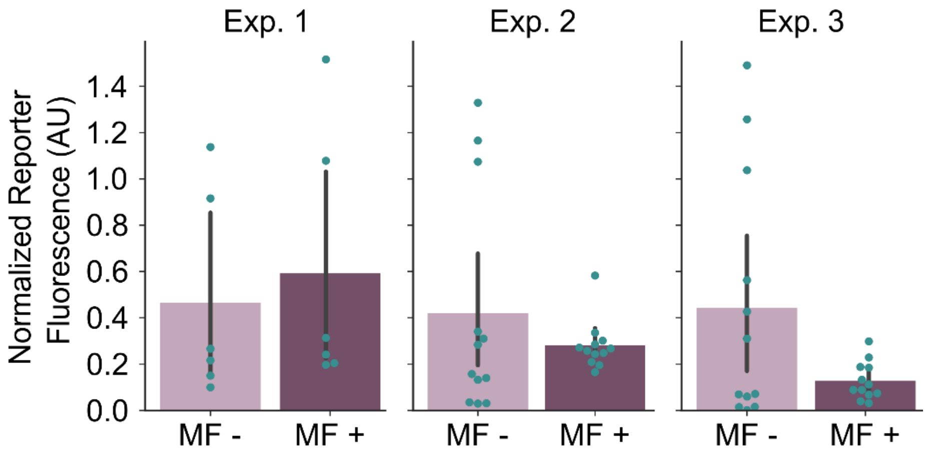


**Supplemental Figure S7. Characterization of the EcR-Cre BR1 control system in stable transgenic lines.** Barplots showing repeated experiments testing function of the EcR-Cre BR1 control system in *A. thaliana* transgenic lines. Across all plots, every dot corresponds to an independent biological replicate.

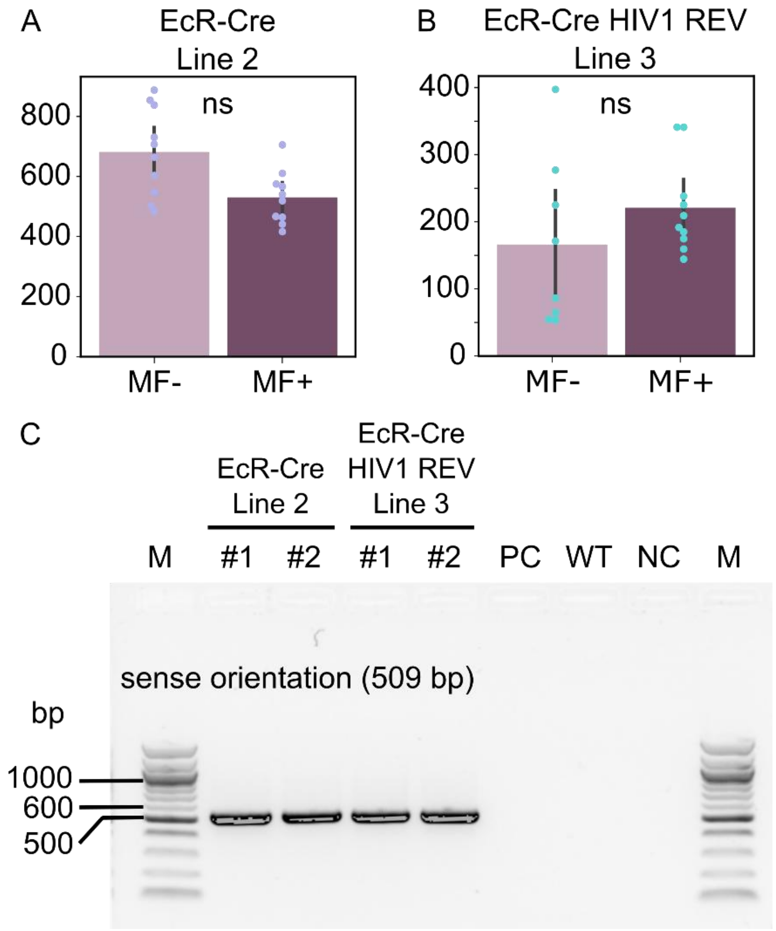


**Supplemental Figure S8. Presence of DNA recombination in the recombinase-based transgenic lines.** A) Barplots representing the change in mScarlet fluorescence after 72 hours of EcR-Cre Line 1 transgenic plants treated with (dark magenta, n=10) and without (light magenta, n=10) MF. B) Barplots representing the change in mScarlet fluorescence after 72 hours of EcR-Cre-HIV1 REV Line 3 transgenic plants treated with (dark magenta, n=10) and without (light magenta, n=8) MF. C) Agarose gel showing PCR amplified products from bulked genomic DNA samples (n=5 seedlings) of the EcR-Cre (Line 1) and EcR-Cre-HIV1 REV (Line 3) control systems. Bands show the presence of the AtUBQ10 promoter in the sense orientation (on' state, 509 bp). Control PCR reactions include a plasmid control containing the T-DNA and the AtUBQ10 promoter in the antisense direction (PC), *A. thaliana* Col-0 WT genomic DNA control (WT), and a reagent negative control (NC). The NEB 100bp DNA Ladder (M) was used for reference. Across all plots, every data point represents a fluorescence measurement from a single seedling. 'ns' represent a non-significant result from a Welch's two sample *t*-test ($p$ < 0.05).

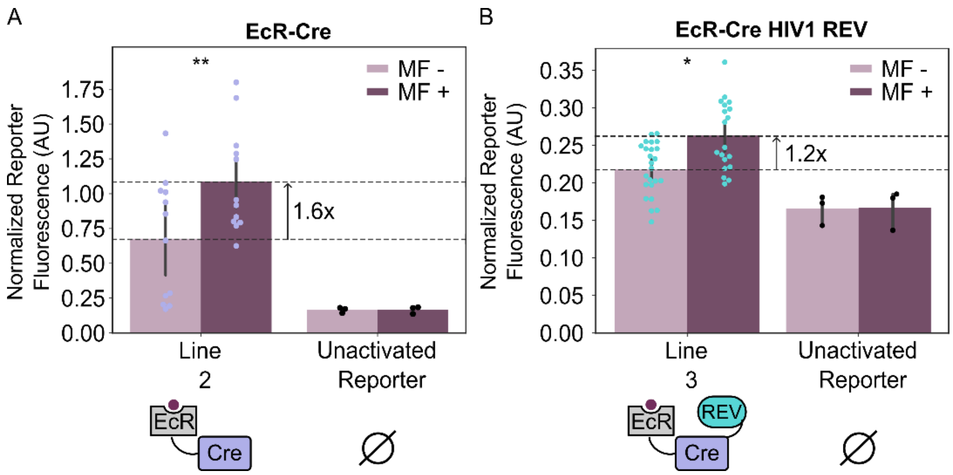


**Supplemental Figure S9. Recovery of the MF-response in the recombinase-based transgenic lines.** A) Barplots representing the normalized reporter fluorescence of EcR-Cre transgenic plants (Line 2) compared to unactivated reporter plants (n=1 independent line, 3 plants per treatment) treated with (dark magenta, MF+, n=19 samples across 4 plants) and without (light magenta, MF-, n=24 samples across 4 plants) MF. B) Barplots representing the normalized reporter fluorescence of EcR-Cre HIV1 REV transgenic plants (Line 3) compared to unactivated reporter plants (n=1 independent line, 3 plants per treatment) treated with (dark magenta, MF+, n=27 samples across 5 plants) and without (light magenta, MF-, n=24 samples across 4 plants) MF. Across all plots, every dot of the same color corresponds to an individual plant from the same line. Asterisks represent results from a Welch's two sample *t*-test ($p < 0.05$), * corresponds to $p < 0.05$, ** corresponds to $p < 0.005$, and *** corresponds to $p < 0.0005$.

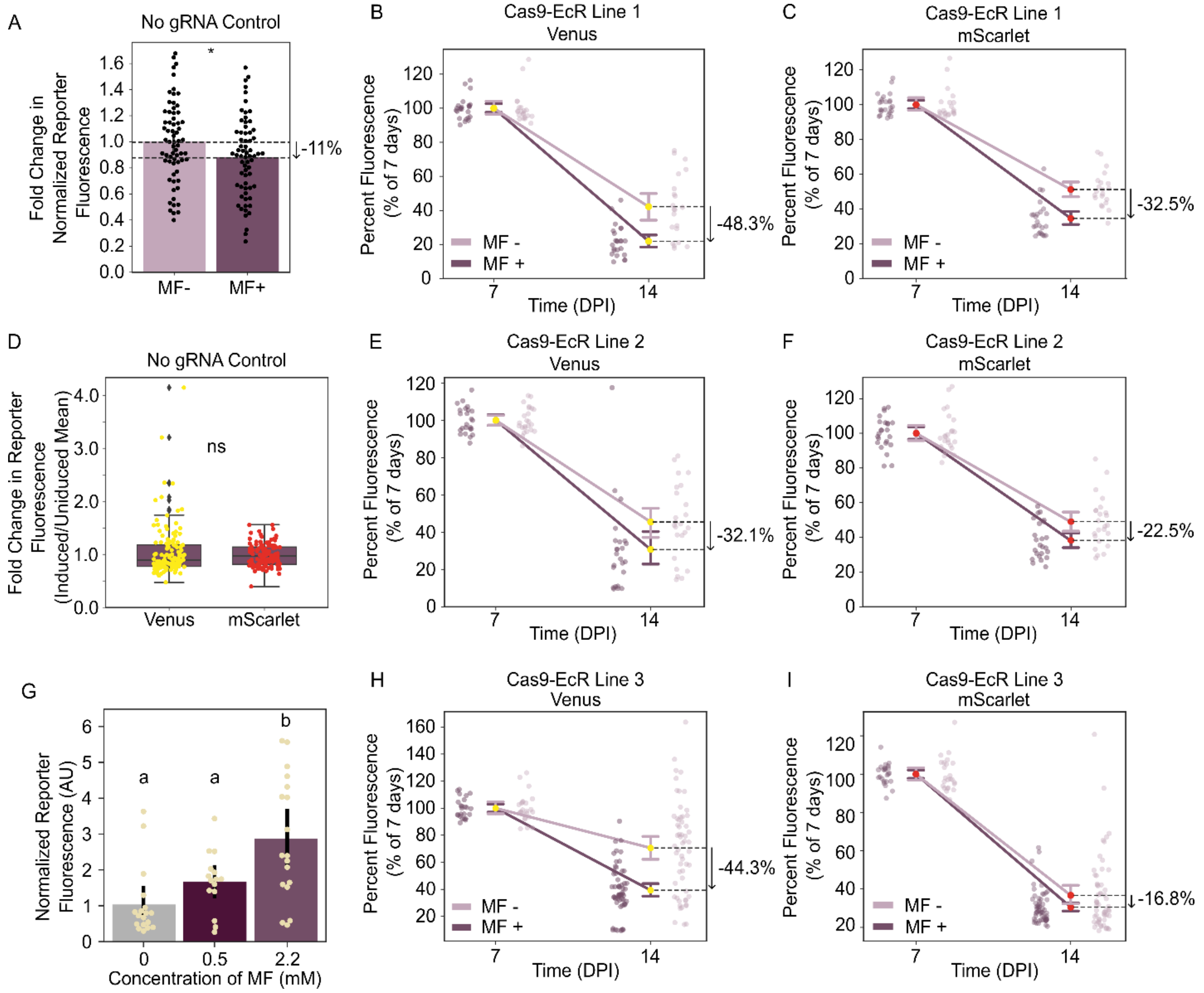


**Supplemental Figure S10. MF-associated effects on gene expression in transgenic lines.** Barplots representing the fold change in normalized reporter fluorescence (induced/uninduced mean) of no gRNA control plants (n=2 independent lines) treated with (dark magenta, n=11 plants per line) and without (light magenta, n=11 plants per line) MF. B, C) Line plots depicting the change in Venus (B) and mScarlet (C) fluorescence in Cas9-EcR Line 1 treated with (dark magenta, n=48 measurements across 8 plants) and without (light magenta, 42 measurements across 8 plants) MF. D) Boxplots representing the fold change in reporter fluorescence (induced/uninduced mean) for Venus and mScarlet fluorophores in the no gRNA population (n=2 independent lines, 11 plants per line) E, F) Line plots depicting the change in Venus (E) and mScarlet (F) fluorescence in Cas9-EcR Line 2 treated with (dark magenta, n=48 measurements across 8 plants) and without (light magenta, 48 measurements across 8 plants) MF. G) Boxplots representing the normalized reporter fluorescence of Cas9-EcR Line treated with 0 (n=15 measurements across 5 plants), 0.5 (n=12 measurements across 4 plants), or 2.2mM (n=15 measurements across 5 plants) MF. H, I) Line plots depicting the change in Venus (H) and mScarlet (I) fluorescence in Cas9-EcR

Line 3 treated with (dark magenta, n=179 measurements across 20 plants) and without (light magenta, 171 measurements across 20 plants) MF. Across all plots, every data point represents a normalized fluorescence measurement from a single leaf disc. Different letters represent statistically significant differences (One-way ANOVA followed by Tukey HSD test, $p < 0.05$). Asterisks represent results from a Welch's two sample *t*-test ($p < 0.05$), * corresponds to $p < 0.05$, ** corresponds to $p < 0.005$, and *** corresponds to $p < 0.0005$. Barplots representing the MF-associated effects on Venus fluorescence at different MF doses.

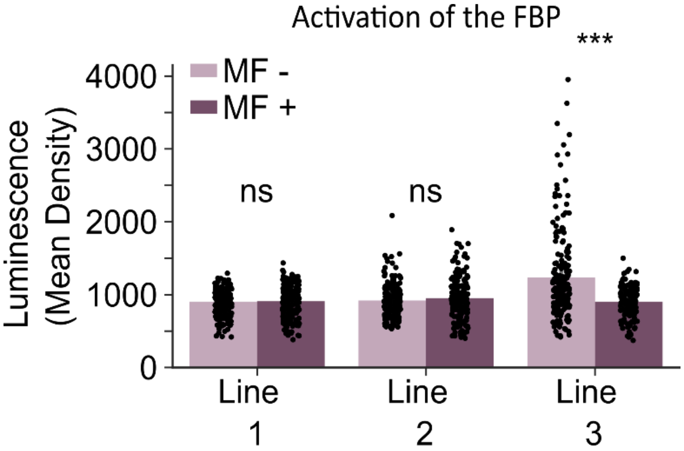


**Supplemental Figure S11. EcR-Cas9 based activation of the FBP in transgenic lines.** Bar plots representing luminescence quantification of three independent *A. thaliana* transgenic lines treated with (dark magenta, n=8 plants per line) and without (light magenta, n=8 plants per line) MF. Every dot corresponds to a luminescence measurement from a single leaf. Asterisks represent results from a Welch's two sample *t*-test ($p < 0.05$), * corresponds to $p < 0.05$, ** corresponds to $p < 0.005$, and *** corresponds to $p < 0.0005$.

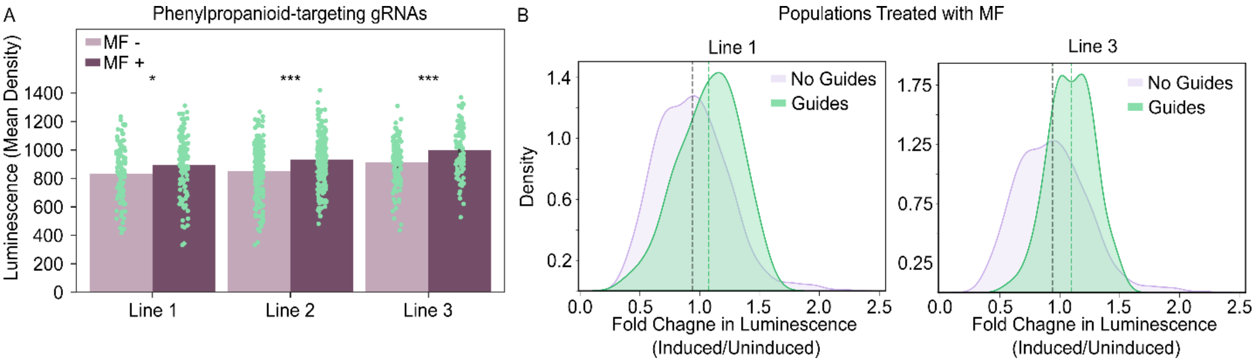


**Supplemental Figure S12. Transcriptionally directing metabolic flux from the phenylpropanoid pathway into the FBP.** A) Barplots representing the luminescence output of three independent transgenic lines that contain the phenylpropanoid targeting gRNAs that were treated with (dark magenta, n=8 plants per line) or without (light magenta, n=8 plants per line) MF. B) Kernel density plots representing distributions in the fold change in luminscence (induced/uninduced) of populations of plants from Line 1 (n=99 measurements across 8 plants) and Line 3 (117 measurements across 8 plants) with phenylpropanoid targeting gRNAs (green) compared to a population of plants without (purple) phenylpropanoid targeting gRNAs (n=578 measurements across 3 independent lines). Asterisks represent results from a Welch's two sample *t*-test ($p < 0.05$), * corresponds to $p < 0.05$, ** corresponds to $p < 0.005$, and *** corresponds to $p < 0.0005$.

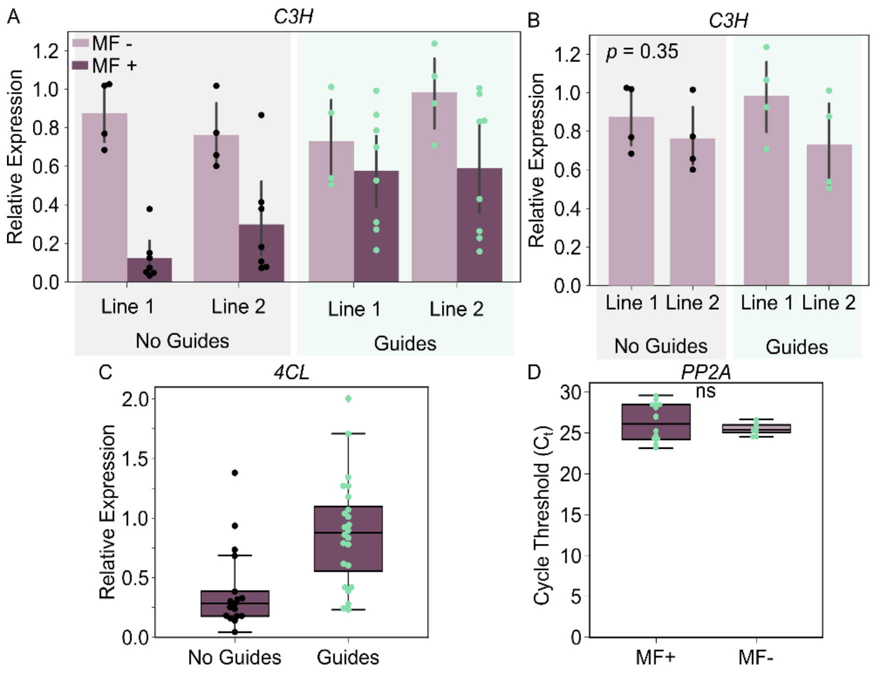


**Supplemental Figure S13. MF-associated effects on phenylpropanoid pathway gene expression.** A, B) Barplots representing relative expression of *C3H* in transgenic lines with (green dots) and without (black dots) phenylpropanoid targeting gRNAs and treated with (dark magenta) and without (light magenta) MF. Each data point represents relative expression of pooled leaf tissue from a single plant C) Relative expression of *4CL* in transgenic plants with (green dots, n=2 independent lines) and without (black dots, n=3 independent lines) phenylpropanoid targeting gRNAs and treated with MF. D) Boxplots representing the cycle threshold values ($C_t$) of *PP2A*, the qPCR housekeeping normalization control from a subset of plants that were treated with (dark magenta, MF+) and without MF (light magenta, MF-). Across all plots, each data point represents relative expression from pooled leaf tissue from a single plant. Asterisks represent results from a Welch's two sample *t*-test ($p < 0.05$), * corresponds to $p < 0.05$, ** corresponds to $p < 0.005$, and *** corresponds to $p < 0.0005$.

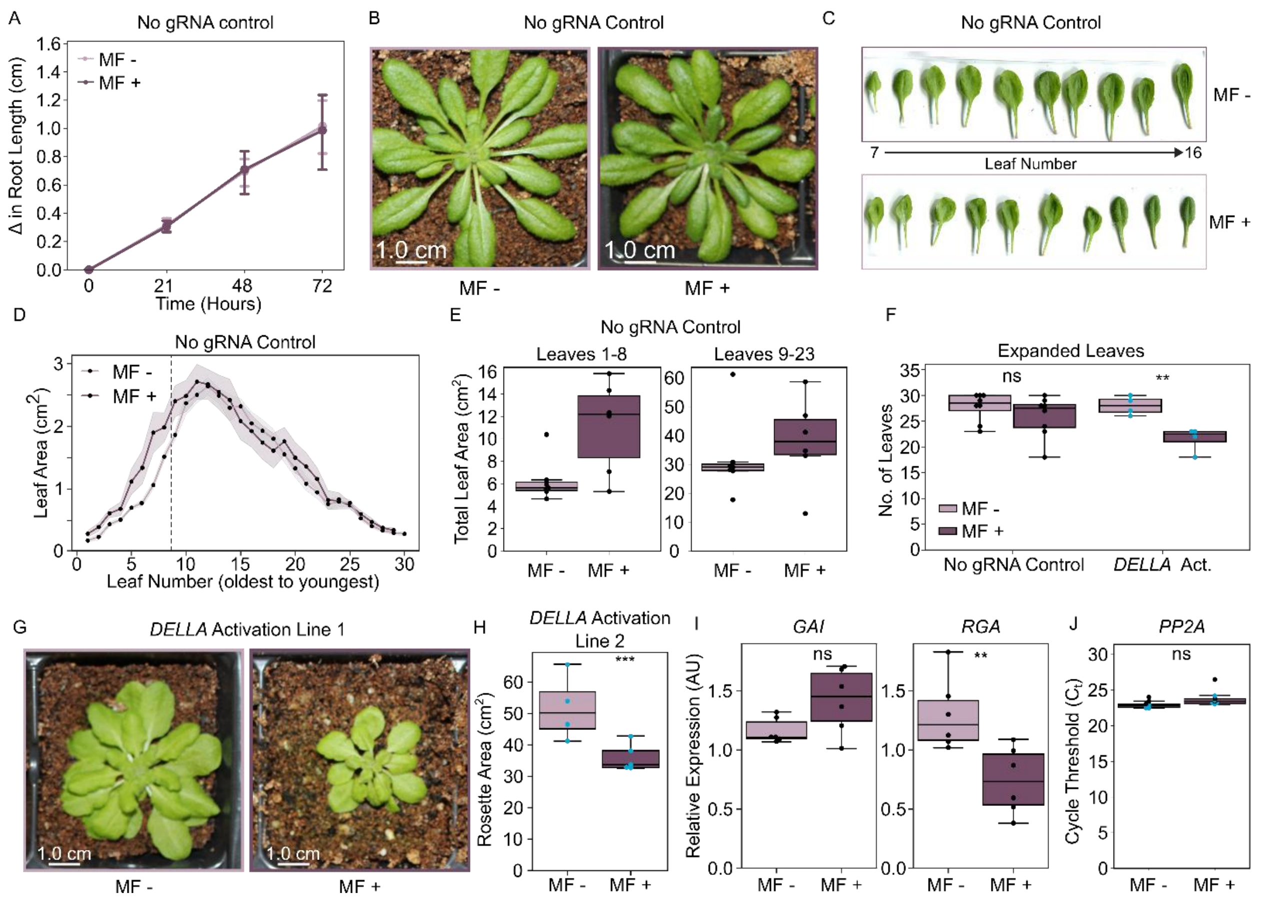


**Supplemental Figure S14. Phenotypic response of Cas9-EcR DELLA activation lines.** A) Line plot representing the change in primary root length of a population of no gRNA control *A. thaliana* independent transgenic lines (n=2 independent lines) treated with (dark magenta, MF+, n=6 or 7 plants) or without (light magenta, MF-, n=6 or 7 plants) MF for 72 hours. B,C) Representative rosette images (B) and leaf images (C) of no gRNA plants treated with (MF+) or without (MF-) MF. D) Line plot representing leaf area across developmental age for plants from the no gRNA population (n=2 independent lines) treated with (dark magenta, n=3 plants per line) or without (light magenta, n=3 plants per line) MF. Each dot represents the mean leaf area across 4 plants. E) Boxplots depicting the total leaf are pre- (left, leaves 1-8, n=2 independent lines, 3 plants per line) and post- (right, leaves 9-23, n=2 independent lines, 3 plants per line) divergence from no gRNA population treated with (dark magenta) or without (light magenta) MF. F) Boxplots depicting the number of expanded leaves for the no gRNA population (n=2 independent lines, 3 plants per line per treatment) and *DELLA* activation Line 1 (4 plants per treatment) of plants treated with (MF+) and without (MF-) MF. G) Representative rosette images of *DELLA* activation plants (Line 1) treated with (MF+) or without (MF-) MF. Each data point represents an individual plant. H) Boxplots representing the rosette areas *DELLA* activation Line 2 treated with (dark magenta, MF+, n= 5 plants) or without (light magenta, MF- n=4 plants) MF. Each data point represents an individual plant. I) Boxplots representing the relative expression of *GAI* and *RGA* in no gRNA control lines (n=2 independent lines) teated with (dark magenta, n=3 plants per line) or without (light magenta, n=3 plants per line) MF. J) Boxplots representing the cycle threshold values ($C_t$) of *PP2A*, the qPCR housekeeping normalization control from a subset of plants that were treated with (dark magenta, MF+) and without MF (light magenta, MF-). Blue dots represent measurements from a single *DELLA* activation Line 1 plant and blakc dots represent a single plant from the no gRNA population. Asterisks represent results from a Welch's two sample *t*-test ($p < 0.05$), * corresponds to $p < 0.05$, ** corresponds to $p < 0.005$, and *** corresponds to $p < 0.0005$.
